# The *in vivo* RNA structurome of the malaria parasite *Plasmodium falciparum*, a protozoan with an A/T-rich transcriptome

**DOI:** 10.1101/2021.04.29.441925

**Authors:** F Dumetz, AJ Enright, J Zhao, CK Kwok, CJ Merrick

## Abstract

*Plasmodium falciparum*, a protozoan parasite and causative agent of human malaria, has one of the most A/T-biased genomes sequenced to date. This may give the genome and the transcriptome unusual structural features. Recent progress in sequencing techniques has made it possible to study the secondary structures of RNA molecules at the transcriptomic level. Thus, in this study we produced the *in vivo* RNA structurome of a protozoan parasite with a highly A/U-biased transcriptome. We showed that it is possible to probe the secondary structures of *P. falciparum* RNA molecules *in vivo* using two different chemical probes, and obtained structures for more than half of all transcripts in the transcriptome. These showed greater stability (lower free energy) than the same structures modelled *in silico*, and structural features appeared to influence translation efficiency and RNA decay. Finally, we compared the *P. falciparum* RNA structurome with the predicted RNA structurome of an A/T-balanced species, *P. knowlesi*, finding a bias towards lower overall transcript stability and more hairpins and multi-stem loops in *P. falciparum*. This unusual protozoan RNA structurome will provide a basis for similar studies in other protozoans and also in other unusual genomes.

## INTRODUCTION

RNA molecules, composed of single strands of four different ribonucleotides, do not adopt a single canonical structure like the double helix of DNA; instead they can fold into complex secondary structures made of loops, hairpins and bulges (Mathews et al. 1999). They serve a variety of roles in the cell (Sharp 2009; Cech and Steitz 2014), including structural roles carried out by transfer and ribosomal RNAs (tRNAs, rRNAs) and protein coding roles for messenger RNAs (mRNAs). These roles are facilitated by the ability of RNA molecules to adopt, and transition between, highly specific secondary structures (Wan et al. 2011; Vandivier et al. 2016). Different secondary structures can impact on the location of the RNA molecules, their metabolism and stability (Sun et al. 2019), their interaction with RNA binding proteins (Casas-Vila et al. 2020), their function and their regulation of protein expression. Two outstanding examples of this are the function-related shape of tRNA (Hasler and Meister 2016) and the translation expression controlling element, the Riboswitch (Wachter 2010).

One of the common strategies to investigate RNA structures *in vivo* is Selective 2′-Hydroxyl Acylation analyzed by Primer Extension (SHAPE) (Wilkinson et al. 2006). SHAPE uses chemicals that modify the 2’ hydroxyl group of flexible nucleotides, in which the 2’ O-adduct will inhibit reverse transcription during cDNA synthesis (Spitale et al. 2013). The pattern of stalling is then detected on a sequencing gel, which determines the identity of all unpaired/flexible bases in the RNA, and hence allows its secondary structure to be inferred. The technicalities of SHAPE only permit the analysis of one transcript at the time, requiring a high amount of starting RNA and potentially more than one sequencing gel to analyse larger transcripts. All these technical limitations rendered impossible the transcriptome-wide study of RNA structures, which is referred to as a structurome (Westhof and Romby 2010). However, the advent of Next Generation Sequencing (NGS) and bio-informatics revolutionised the field of RNA structure and within the past decade, Structure-seq (Ding et al. 2014), and many other related methods (Kwok et al. 2015; Zhao et al. 2019; Rouskin et al. 2014), were developed.

To date, only a few *in vivo* structuromes are available. Many viruses have been studied, probably due to their small number of RNA molecules, such as HIV (Watts et al. 2009; Tomezsko et al. 2020), DENV and ZIKV (Li et al. 2018; Huber et al. 2019), influenza (Simon et al. 2019) and more recently SARS-CoV-2 (Manfredonia et al. 2020; Sun et al. 2021; Ziv et al. 2020; Huston et al. 2021; Ling Yang et al. 2021; Tammy C. T. Lan, Matthew F. Allan, Lauren E. Malsick, Stuti Khandwala, Sherry S. Y. Nyeo, Yu Sun, Junjie U. Guo, Mark Bathe, Anthony Griffiths et al. 2021). Regarding eukaryotes, structuromes were established for model organisms like the plants *Arabidopsis thaliana* (Ding et al. 2014) and *Oryza sativa* (rice) (Deng et al. 2018), the yeast *Saccharomyces cerevisiae* (Rouskin et al. 2014), the bacteria *Escherichia coli* (Guo and Bartel 2016) and amongst mammals, mouse embryonic stem cells (Incarnato et al. 2014; Guo and Bartel 2016) and human cell lines (Sun et al. 2019; Rouskin et al. 2014; Wu and Bartel 2017; Guo and Bartel 2016). These have revealed fundamental conserved features across the landscape of RNA structures.

In this paper we have established the *in vivo* structurome of a protozoan, *Plasmodium falciparum*. *P. falciparum* is one of the etiological agents of malaria, the most widely lethal human parasitic disease in the world (WHO 2018). Little is known about RNA structures in *P. falciparum*, yet the parasites offer many opportunities to understand the roles of RNA structures in eukaryotic cells. Firstly, the natural life cycle of the parasite oscillates from a warm-blooded host to an insect vector, meaning that the same organism is exposed to very different temperatures and metabolic environments. This allows the study in natural conditions of the effect of the environment and temperature upon the RNA structurome. Indeed, it has been shown that heat stress can affect the structuromes of plants (Su et al. 2018) and bacteria (Righetti et al. 2016; Twittenhoff et al. 2020), which can likewise experience temperature extremes in nature. Furthermore, one intriguing piece of evidence concerning rRNAs suggests that temperature-responsive RNA structures are also important in *Plasmodium* life cycles. *Plasmodium* does not encode rRNAs in conventional tandem arrays, but rather in several isolated units which are transcribed independently in either mosquito stages (S-type) or mammalian stages (A-type) (Waters et al. 1989; Gunderson et al. 1987). rRNA transcription apparently switches in response to differential temperature and nutrient availability in the two hosts (Fang and McCutchan 2002; Fang et al. 2004). The two types of rRNA were predicted, based on their sequences, to have different structures (John Rogers et al. 1996) and they do perform differently when complemented into *S. cerevisiae* (Velichutina et al. 1998), so there is clear potential for them to influence translation in insects versus humans. Thus, *Plasmodium* may have evolved variant RNA structures as a key element in its life cycle transitions.

Secondly, while all the previously cited organisms have a more balanced ratio of A/T and G/C in their genomes, *P. falciparum* is 80.6% A/T-rich (Gardner et al. 2002): one of the most biased genome compositions ever to be sequenced. This offers a critical point of view when studying genome evolution, and it may well lead to unusual RNA structures.

This RNA structurome of *P. falciparum* sets out intends to investigate the relationship between RNA shapes and genome composition, as well as transcription efficiency, in the asexual erythrocytic cycle of this important human parasite.

## RESULTS

### Assessment of NAI and DMS as RNA structure probing chemicals for *P. falciparum* inside the erythrocyte host cell

We conducted *in vivo* structure-seq on *Plasmodium falciparum*: an organism with a >80% AT-rich genome, and one that lives entirely inside another host cell. It was necessary first to test and optimise the parameters for *in vivo* RNA probing in this unusual organism. We used two different probing agents with different nucleotide reactivity: NAI (2-methylnicotinic acid imidazolide) and DMS (dimethyl sulfate). NAI reacts with all four unpaired nucleotides while DMS selectively reacts with unpaired adenosine and cytosine. Neither of these chemicals had been used in *P. falciparum* before. In order to check if the chemicals could penetrate inside the parasite inside the host erythrocyte, we exposed an asynchronous culture of *P. falciparum* to NAI or DMS. After RNA extraction we performed a primer extension with a Cy5-conjugated primer targeting the 5.8S rRNA transcript. When a nucleotide has been modified by one of the chemicals, meaning that the nucleotide is unpaired, the reverse transcriptase will stall one nucleotide beforehand, stopping the elongation reaction (Fig. 1A). This can be seen on a denaturing polyacrylamide sequencing gel (Fig. 1B, Supplemental Fig. 1). Reading the sequence confirmed that it was identical to the PF3D7_0531800 gene sequence available on PlasmoDB (Aurrecoechea et al. 2009). In NAI and DMS probed parasites, base reactivity was seen with all four nucleotides after NAI-probing, and with only adenosine and cytosine after DMS-probing. (Modification of adenosine and cytosine was not, however, identical at all positions with DMS and NAI, because these chemical probes react with different moieties of the RNA nucleotides, and with different efficiencies.) This indicated that both chemicals could penetrate into *P. falciparum*, and our probing strategy could investigate the RNA structures in *P. falciparum*.

**Figure 1:**
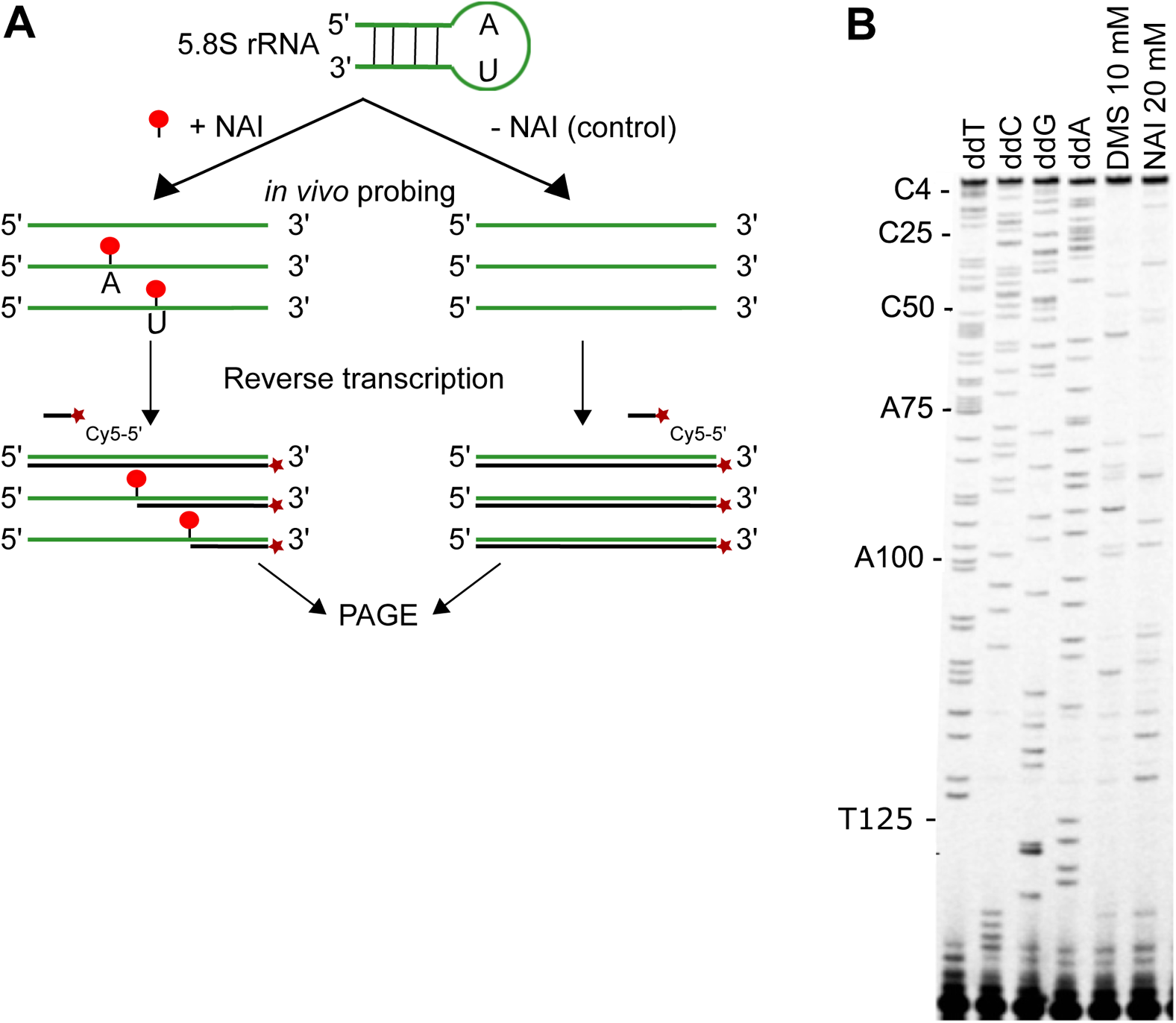
*In vivo* probing of RNA structures in *P. falciparum*. **(A)** Schematic of RNA structure probing. An asynchronous parasite culture was split into two, one exposed to NAI (+NAI) and the other exposed to DMSO as control (−NAI). Reverse transcription (RT) was performed using a 5’-Cy5 conjugated primer (dark red star) specific for the 5.8S rRNA of *P. falciparum* (in green). During RT, nucleotides that have been modified by NAI in the +NAI samples (red dots) will stall the reverse transcription, interrupting the elongation process (black line). Finally, elongation cDNA products are resolved on a PAGE gel. **(B)** Sequencing gel showing the 5.8S rRNA gene of *P. falciparum.* Lanes 1 to 4 are sequencing lanes where one of the four nucleotides in the reaction mix has been switched with the dideoxynucleotide, provoking RT-termination. Lanes 5 and 6 represent respectively DMS and NAI modifications on 5.8S rRNA.

To prepare the structurome, we then extracted the total RNA from two independent asynchronous parasite cultures treated with NAI, DMS, or DMSO only, as in figure 1, and used poly(T) magnetic beads to enrich poly(A) RNA. The enriched poly(A) fraction represented 2-3.4% of the total RNA, which is commensurate with the literature for eukaryotic cells (Westermann et al. 2012). The enriched fractions were then used to generate NGS libraries (Fig. 2A). The sequencing outputs were processed using *StructureFold2* and *RNAStructure* for experimental reactivity constrained folding (Ritchey et al. 2017) to determine the secondary structure of the RNA molecules. Each replicate for each chemical treatment was analysed separately. After preliminary analysis, because the distribution of structure differences between replicates was marginal and the correlation between replicates and pooled samples was good (Supplemental Fig. 2), reads from both replicates were merged for subsequent analysis. To ensure structure calls of a high quality, mappings were filtered such that there were no more than 3 mismatches/indels from the reference and the first 5’ base was matching. Coverage filtering also required transcripts with complete coverage between samples (i.e. meeting the *StructureFold2* default filtering threshold of an average coverage >1.0 per nucleotide throughout the transcript (Supplemental Fig. 3). Structural information was more comprehensive (and agreement between replicates was also better), for the NAI-probed datasets because all four unpaired nucleotides were detected. Subsequent analysis was therefore focused on the NAI dataset as the primary *in vivo* structurome.

**Figure 2:**
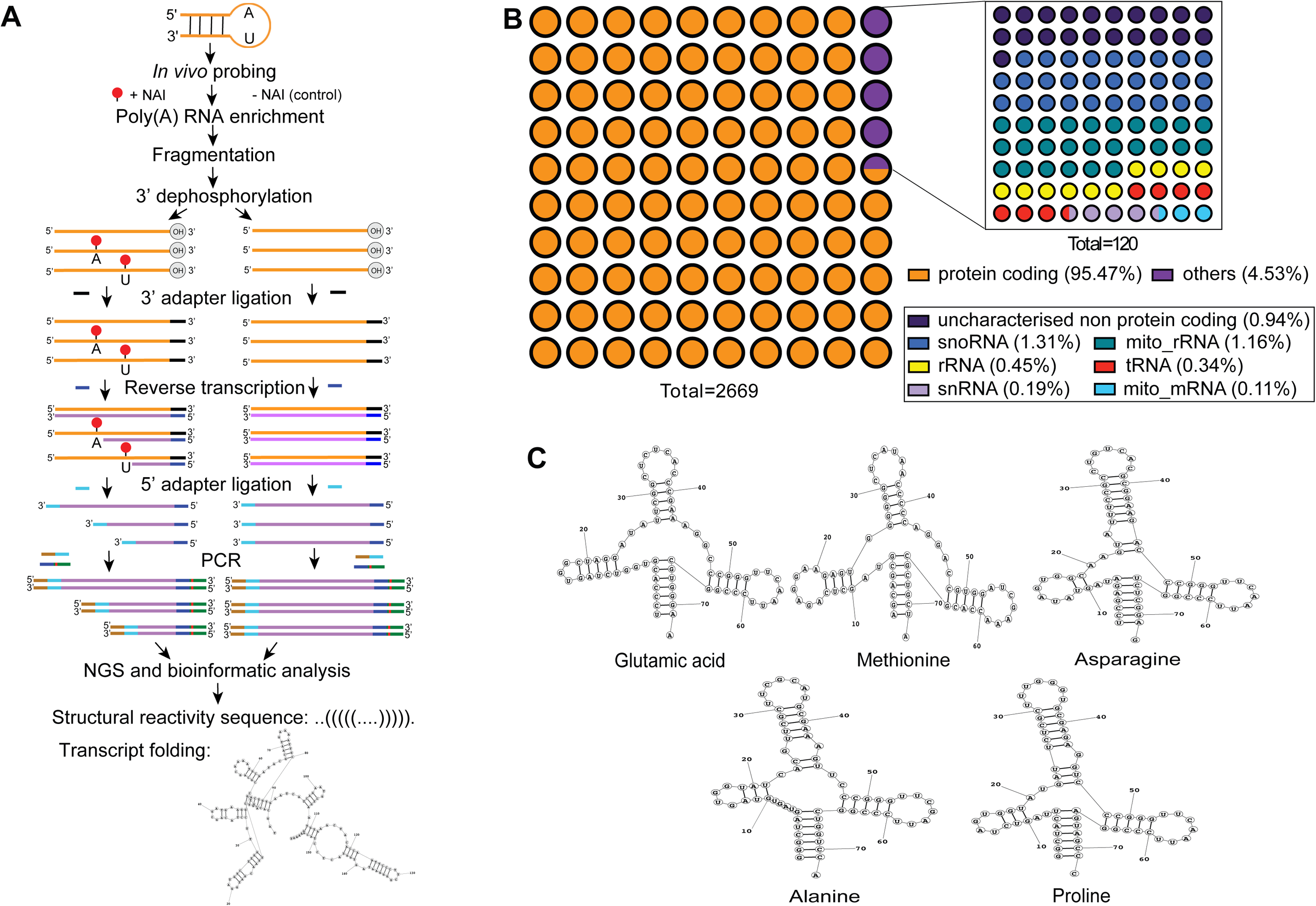
RNA structurome of *P. falciparum* using Structure-seq. **(A)** Schematic showing the Structure-seq pipeline. An asynchronous parasite culture was split into two, one exposed to NAI (+NAI) and the other exposed to DMSO as control (−NAI). Polyadenylated-enriched RNA fractions were then used to prepare the sequencing libraries in parallel. Fragmentation was performed to generate an average length (~250 nt) of RNA for sequencing followed by 3’ dephosphorylation, which replaces the 2’3’ cyclic phosphate group with a 3’-OH group for 3’ adapter ligation. Then a 3’ adapter (black line) was ligated to the 3’-OH group of dephosphorylated RNA. Next, RT of RNA molecules (orange line) was carried out using a designed reverse primer (dark blue line) at the 5’ end. During RT, nucleotides that had been modified by NAI in the +NAI samples (red dots) would stall the reverse transcription, interrupting the elongation process (pink line). Later, a 5’ adaptor (light blue line) was added by ssDNA ligation at the 3’ end of the newly synthesised cDNA strand (pink line). Finally, forward primers (brown+light blue line) and reverse primers (dark blue+red+green line) with different indexes were added by PCR for NGS and bioinformatic analysis. Reads were aligned to the PF3D7 reference genome and subsequently analysed with *StructureFold2* to determine the structural reactivity sequence (transcribed as a dot/bracket sequence denoting paired/unpaired nucleotides), and with *RNAStructure* to predict the folding. **(B)** Representation of the diversity of the dataset regarding RNA families. **(C)** Structure-seq derived structures of glutamic acid (PF3D7_0411600), methionine (PF3D7_1339100), asparagine (PF3D7_0714700), alanine (PF3D7_0620800) and proline (PF3D7_1339200) tRNAs, all of which are similar to canonical tRNA structures.

### Structure of the NAI-probed dataset

After filtering for low-coverage genes, the NAI structurome covered 51.23% of the total transcriptome of *P. falciparum* (2699 transcripts detected out of 5268 total). We first looked at the representation of the various families of RNA molecules. Within our dataset 95.5% of the transcripts were protein coding transcripts, 1.27% corresponded to mitochondrial transcripts (1.16% mitochondrial rRNA, 0.11% mitochondrial mRNA) and 0.94% were uncharacterised non-protein coding transcripts. Other types of non-coding RNA were also detected, 1.31% of snoRNA, 0.19% of snRNA and 0.34% of tRNA (Fig. 2B). Within the protein coding transcripts, nine were encoded in the apicoplast, a subcellular organelle that is unique to apicomplexans and represents a relic chloroplast (Supplemental table 1).

As well as the different families of transcripts, we detected 114 transcripts with splice variants (Supplemental table 2). However, since we used a short-read technology to sequence the library, it was impossible to associate all the reads within a variant transcript to one variant. Nevertheless, this overall analysis indicates that our *in vivo* chemical probing, RNA extraction and polyA selection were robust because all the different RNA families in *P. falciparum* were detected, with a strong bias towards mRNAs, and if a long-read sequencing technique had been used, even the detection of transcript variants would likely have been possible.

Some RNA families have their function dictated by their structure. This is especially the case of tRNA. Within our dataset 9 tRNAs were identified. Using *RNAStructure* we were able to call their structures and 5 were similar to canonical eukaryotic tRNA molecules. Those tRNAs were for glutamic acid (PF3D7_0411600, PF3D7_0527700), methionine (PF3D7_1339100), asparagine (PF3D7_0714700), alanine (PF3D7_0620800) and proline (PF3D7_1339200) (Fig. 2C). In the other 3 tRNAs, for valine (PF3D7_0312600), serine (PF3D7_0410100) and leucine (PF3D7_0620900), the lowest-free-energy structures predicted by *RNAStructure* were not canonical (Supplemental Fig. 4). For some but not all of these, alternative folding with less favourable energy produced canonical structures, albeit less compatible with the data defining free/flexible bases (e.g. Supplemental Fig. 4A). It is theoretically possible that dynamic structural variants of some tRNAs exist *in vivo*, although it is unclear why some tRNAs would differ from others in the dataset. Importantly, however, tRNA structure is also dependent on nucleotide base modification such as m^2^A, m^2^G, pseudouridine, etc. [46]. Those modifications have not been assessed in this study, but it is possible that they would influence the apparently-noncanonical tRNA structures.

In general, opportunities to ‘verify’ the structurome were limited because of the lack of *Plasmodium* RNA structures obtained independently, by orthogonal experimental methods. The canonical structural model of a blood-stage *P. falciparum* rRNA is available from the Comparative RNA Web (CRW) site and this was used to map base reactivities obtained *in vivo* (Supplemental Fig. 5). The result demonstrates visually the A/C reactivity of DMS versus the all-base reactivity of NAI, and makes it clear that many bases modelled as unpaired are nevertheless not highly reactive *in vivo*. Other factors such as protein binding and base modification probably affect their reactivity, emphasising the over-simplification imposed by *in silico* modelling and the extra level of information obtained by *in vivo* structural probing.

### Divergence between *in vivo* RNA structures and structures modelled *in silico*

Using the *RNAStructure* algorithm, RNA structures can be predicted from primary sequence alone, or they can be predicted with the addition of base-pairing information gained experimentally. We generated both types of structure, transcriptome-wide, taking only the lowest-free-energy structure in all cases where the algorithm generated several alternatives. This was simply because, in the absence of other constraints, the lowest-free-energy structure is the most likely to form.

Firstly, we sought to discover how much these two types of structures would differ in *P. falciparum*, i.e. how much new information was gained by performing the *in vivo* structurome? We calculated the ratio of bases that differed (i.e. that were paired versus unpaired) between each *in-silico-*predicted and probing-informed structure, and termed this ratio the ‘divergence’. Most of the non-protein coding transcripts had relatively low divergence, meaning that they were minimally informed by probing (Table 2). This was expected for highly conserved, stereotypical structures like rRNAs and tRNAs – although the divergence for mitochondrial rRNAs, which are encoded in a highly fragmentary fashion in the *P. falciparum* mitochondrial genome, was notably greater than for nuclear-encoded rRNAs. By contrast, the divergence of protein coding transcripts was much higher: a median of 0.7067 for the nuclear-encoded mRNAs and 0.8117 for mitochondrial mRNAs (Table 2).

**Table 2:** Distribution of ‘divergence’ normalised by transcript length between base pairing determined by Structure-seq and *in silico*.

|  | Protein coding | mito_m RNA | Uncharacterized non protein coding | snoRNA | tRNA | rRNA | snRNA | mito_rRNA |
| --- | --- | --- | --- | --- | --- | --- | --- | --- |
| # transcript | 2549 | 3 | 25 | 35 | 9 | 8 | 5 | 36 |
| Median | 0.7067 | 0.8117 | 0.5260 | 0.4414 | 0.1667 | 0.2204 | 0.4630 | 0.5439 |
| Minimum | 0.04408 | 0.7480 | 0.07407 | 0.07500 | 0.0556 | 0.0202 | 0.1880 | 0.000 |
| 25% Percentile | 0.5995 | 0.7480 | 0.3428 | 0.2639 | 0.0972 | 0.0616 | 0.1916 | 0.3471 |
| 75% Percentile | 0.7874 | 0.8880 | 0.6914 | 0.6341 | 0.6001 | 0.3529 | 0.5636 | 0.7446 |
| Maximum | 1.033 | 0.8880 | 0.9083 | 0.9176 | 0.7317 | 0.4706 | 0.6566 | 1.050 |

Since structure and function in non-protein coding transcripts, like tRNAs, are deeply intertwined, we then sought any relationship between folding complexity and biological function. Using the ‘divergence’ as a metric, we split the protein coding transcripts into 3 tiers: minimal value to 25^th^ percentile, 25^th^ to 75^th^ percentile and 75^th^ percentile to the highest value (Table 2), The list of transcripts was then submitted for gene ontology enrichment and using Revigo (Supek et al. 2011) we obtained broad families of GO terms and networks of terms belonging to a similar biological pathway. In tier 1, with the lowest ‘divergence’, we observed an enrichment of 25 terms, with the strongest enrichment in terms related to ribosomal functions. This was expected, since ribosomal RNA structures are highly conserved and are likely to be predicted accurately *in silico* with little alteration in *P. falciparum*. Other terms such as ‘macromolecular complex” were strongly enriched in tier 1, but they were not part of any network (Fig 3A, Supplemental table 3-1). Twenty-one terms were found in tier 2. Two terms were strongly enriched: ‘translation’ and ‘cellular amide metabolism’; however, these stand alone. Two networks could be created: a large one around ‘protein’ and ‘protein regulation’ (composed only of lowly enriched words) and a second network restricted to two terms around RNA splicing (Fig 3B, Supplemental table 3-2). In tier 3, with the lowest level of identity between *in vivo* structures and *in silico* predictions, only 11 terms were enriched and 3 networks were made, the largest composed of 7 terms strongly enriched around different kinds of metabolism and the others around transport and complex molecule biosynthesis (Fig 3C, Supplemental table 3-3).

**Figure 3:**
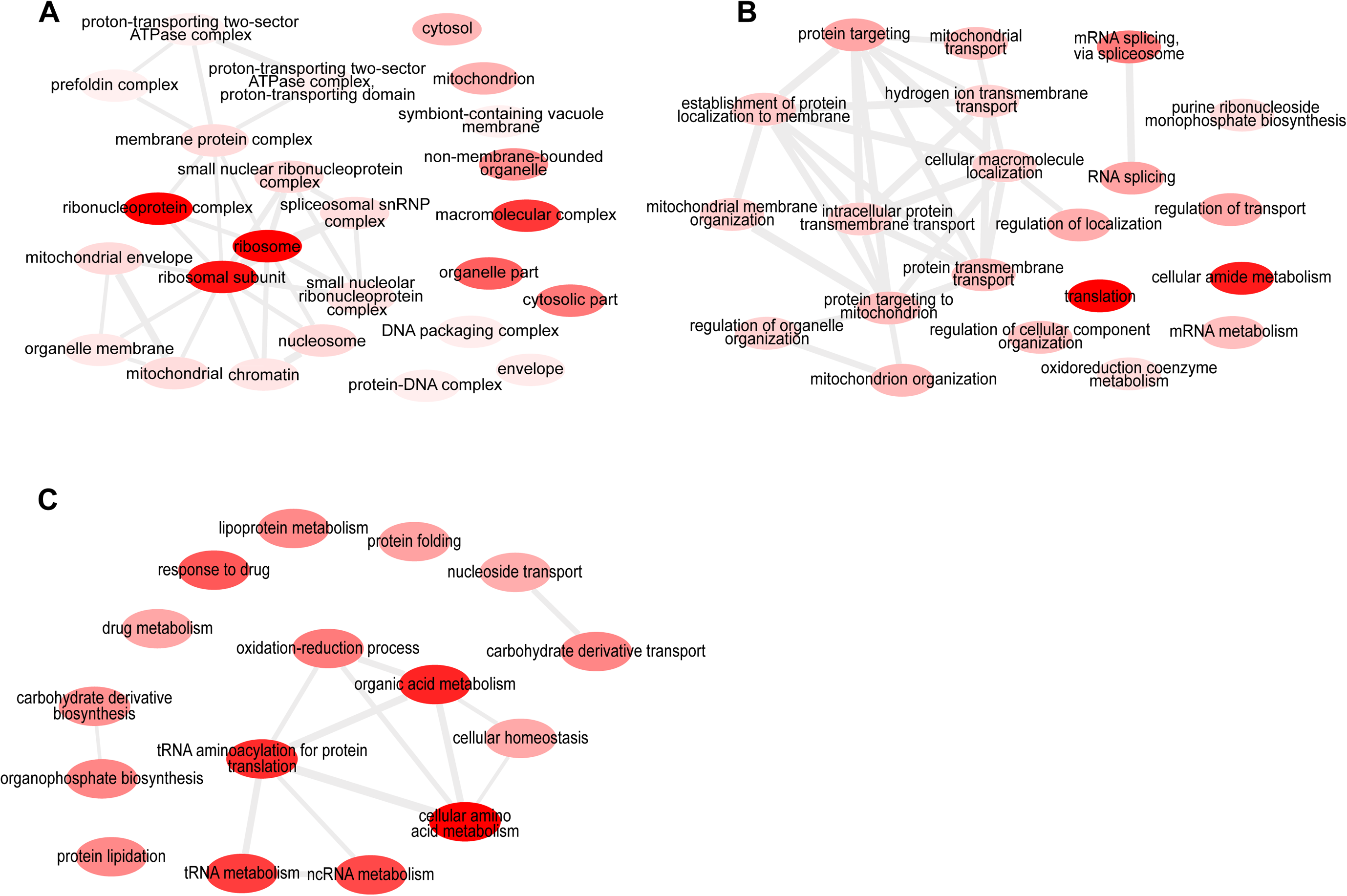
GO terms enrichment networks for protein coding transcripts, tier 1 to 3. All networks represent the GO enrichment provided by REVIGO for **(A)** tier 1, **(B)** tier 2 and **(C)** tier 3. The brighter the red, the more enriched the term is. Detailed values are available in Supplemental table 3.

Secondly, we assessed the value of performing Structure-seq instead of just *in silico* prediction for modelling the shape of RNA molecules. We used ViennaFold on the *P. falciparum* transcriptome to fold the structures with or without reactivity data. We first aligned the dot/bracket sequences obtained from *in silico* folding with the Structure-seq-determined sequence of the U2 snRNA (PF3D7_1137000). It showed a difference in the pairing of 74 nucleotides out of 198, meaning a 37.4% discrepancy in the structure determined *in silico* compared to *in vivo* (Fig. 4A).

**Figure 4:**
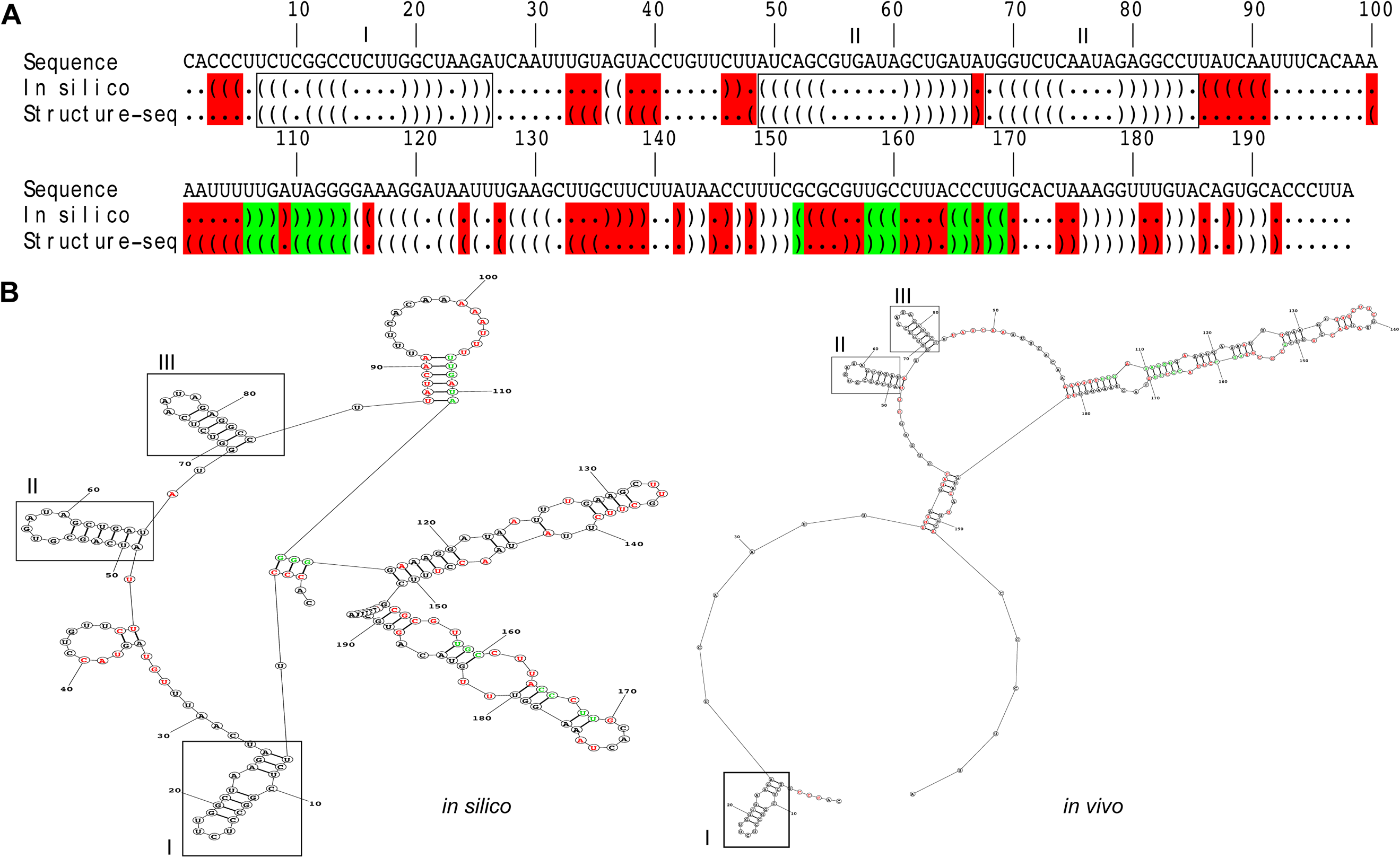
Difference in U2 snRNA structure determined by Structure-seq versus *in silico* prediction. **(A)** U2 snRNA (PF3D7_1137000) reactivity sequence alignment, comparing *in vitro* probing using NAI and *in silico* prediction using the same constraint parameters. The first line shows the nucleotide sequence, the second line shows the *in silico* predicted dot/bracket sequence and the third line, the Structure-seq dot/bracket sequence extracted from reactivity measurement. A dot shows a nucleotide not paired with any other nucleotide. An open bracket shows a nucleotide paired with another and starting a structural motif, while a closed bracket shows a nucleotide paired with another but finishing a structural motif. Red-highlighted dot or bracket indicates a change of pairing between the NAI and *in silico* predicted structural sequence. A green-highlighted bracket indicates a change in motif orientation between the NAI and *in silico* predicted structural sequence. Boxes with roman numerals indicate the conserved structures between the two shapes. **(B)** U2 snRNA structure predicted *in silico* from RNA sequence alone, on the left, and with the addition of reactivity data on the right. The red and green colouring have the same meaning as in (A).

Structurally, the difference between the two transcripts is significant and only 3 loops are common to both structures (Fig. 4B). More interestingly, an RNA-appropriate measurement of Minimum Free Energy, MFEden (Trotta 2014), revealed that the Structure-seq U2 snRNA structure had a lower MFEden (−24.76 versus 15.50).

Extending this to the whole dataset, we observed a similar effect in each RNA category (Table 3). Thus, structures called from Structure-seq are more likely to form and are more stable than the ones predicted only using an *in silico* prediction.

**Table 3:** MFEden average for every type of RNA molecule.

|  | protein<br>coding | mito_<br>mRNA | uncharacterised<br>non protein<br>coding | snoRNA | tRNA | rRNA | snRNA | mito_<br>rRNA |
| --- | --- | --- | --- | --- | --- | --- | --- | --- |
| Structure-seq | -20.5 | -27.0 | -29.8 | -29.6 | -63 | -31.1 | -37.9 | -25 |
| *in silico* | 14.1 | 10.3 | 10.7 | 10.2 | -8.8 | 3.2 | 10.7 | 2.7 |

All MFE<sub>den</sub> values per transcript are in Supplemental table 4.

### Effect of A/T bias in the structurome of *P. falciparum* by comparison with an A/T-balanced *Plasmodium* species

The *Plasmodium* genus offers the possibility to compare very different genome compositions. *P. falciparum* is ~80% A/T rich while *P. knowlesi*, a macaque *Plasmodium* species which also infects humans, is 62.5% A/T rich (Pain et al. 2008). In order to assess whether genome base composition could affect the structurome, we compared the *in silico* folding of *P. falciparum* and *P. knowlesi* structuromes. Firstly, we assessed the stability of the two predicted structuromes by comparing the average free energy (MFEden (Trotta 2014)) of all folded structures: *P. falciparum* had a significantly higher MFEden than *P. knowlesi* (respectively 14.03 and 7.1, t-test *p*-value <0.0001). Secondly, we looked at the occurrence of various structures by comparing the length-normalised number of nucleotides participating in a structure (e.g. a hairpin or bulge) in each *P. falciparum* transcript with their *P. knowlesi* orthologs. The structurome of *P. falciparum* had significantly more nucleotides involved in hairpins and multi-stem loops than the structurome of *P. knowlesi* (t-test, *p*-value <0.0001), whereas *P. knowlesi* had significantly more nucleotides involved in stems and bulges (t-test, *p*-value <0.0001) (Supplemental table 5).

### RNA structures affect translation efficiency in a stage specific manner

Figure 2 shows that various families of RNAs display very conserved structures that are related to their particular functions. This is clearly true for structural RNAs (tRNAs, rRNAs, etc.), but it may also be true for mRNAs, whose main function is to be translated into proteins. The stability of the transcript and the efficiency of its translation could both be affected by mRNA structure. Therefore, if mRNA structures can vary with varying cellular conditions, transcript and protein abundance may both vary accordingly.

We first explored the concept that transcript abundance could vary with cellular conditions due to changes in mRNA structure. For this, we used published microarray and transcriptomic datasets from *P. falciparum* under physiological stresses like hyperoxia (Torrentino-Madamet et al. 2011), or drug exposure like chloroquine treatment (Untaroiu et al. 2019). These datasets reveal which transcripts are upregulated and downregulated in each condition. We compared the most differentially expressed genes in each condition with the structurome dataset, to see if they were particularly structured or unstructured. Neither the overall degree of ‘divergence’ nor the representation of diverse structures (the proportion of hairpins, loops, etc), correlated with differential expression under stress (data not shown).

Secondly, we looked at the effect of RNA structures on translation efficiency (TE). To establish the translation efficiency for each transcript in our dataset, we extracted from previously published work (Caro et al. 2014) the average level of ribosome attachment per transcript, determined by Ribo-seq. We obtained TE for the transcriptome of rings, early and late trophozoites and schizonts. Applying a linear regression model, we compared TE with transcript structuredness (‘NAI ratio’, i.e. the ratio of paired to unpaired bases) and also the proportion of different motifs such as stems, hairpins, multi-loop stems and bulges (Fig. 5). We observed no correlation between the transcript-length-normalised ‘structuredness’ and TE in any of the four stages (Fig. 5A). Therefore, there was no overall relationship between the proportion of paired bases in a transcript and the efficiency of its translation. However, breaking down the different possible structures (Fig. 5B-E) we observed a positive correlation of TE with the presence of stems and a negative correlation between TE and hairpins, multi-loop stems and bulges. The correlations were only seen in the more mature parasite stages, late trophozoites and schizonts.

**Figure 5:**
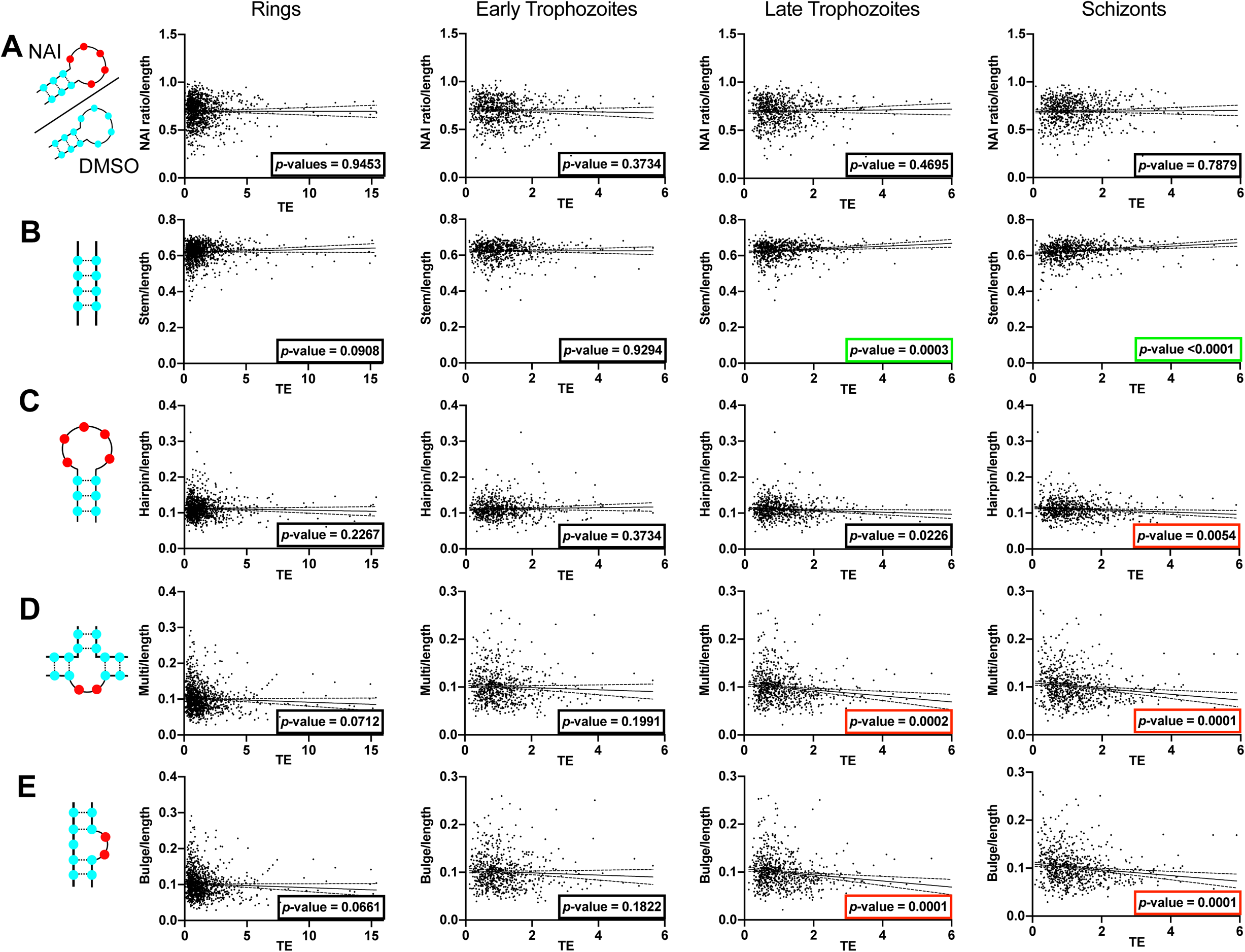
Impact of various RNA motifs on transcription efficiency across the life cycle of *P. falciparum*. The columns represent the different erythrocytic life stages of *P. falciparum* and the rows represent the correlation between transcription efficiency and **(A)** the transcript structuredness normalised by the transcript length, **(B)** the number of stems, **(C)** the number of hairpins, **(D)** the number of multi-stem loops and **(E)** the number of bulges. Statistically significant positive correlations have a *p*-value framed in green and statistically significant negative correlations have a *p*-value framed in red.

### RNA decay is mediated by the same structures across the cell cycle

The final critical part of an RNA molecule’s life inside the cell is its rate of decay. Following a similar reasoning as for translation efficiency, we extracted RNA decay values from data published by Painter *et al*. (Painter et al. 2018) and tested for any correlation between RNA decay and RNA secondary structures in the different cell cycle stages (rings, early and late trophozoites and schizonts). For this, we applied a linear regression model to compare RNA decay (the higher the value, the less stable the molecule is) to the number of different structures per transcript. We observed no correlation between RNA decay and RNA secondary structures involving high amounts of base pairing: stems and hairpins (Fig. 6A, B). However, when nucleotides were more exposed, as in bulges and multi-stem loops (Fig. 6C, D) we observed correlations between RNA decay and the presence of those structures in all stages (except for multi-stem loops in ring-stage) (Fig. 6C).

**Figure 6:**
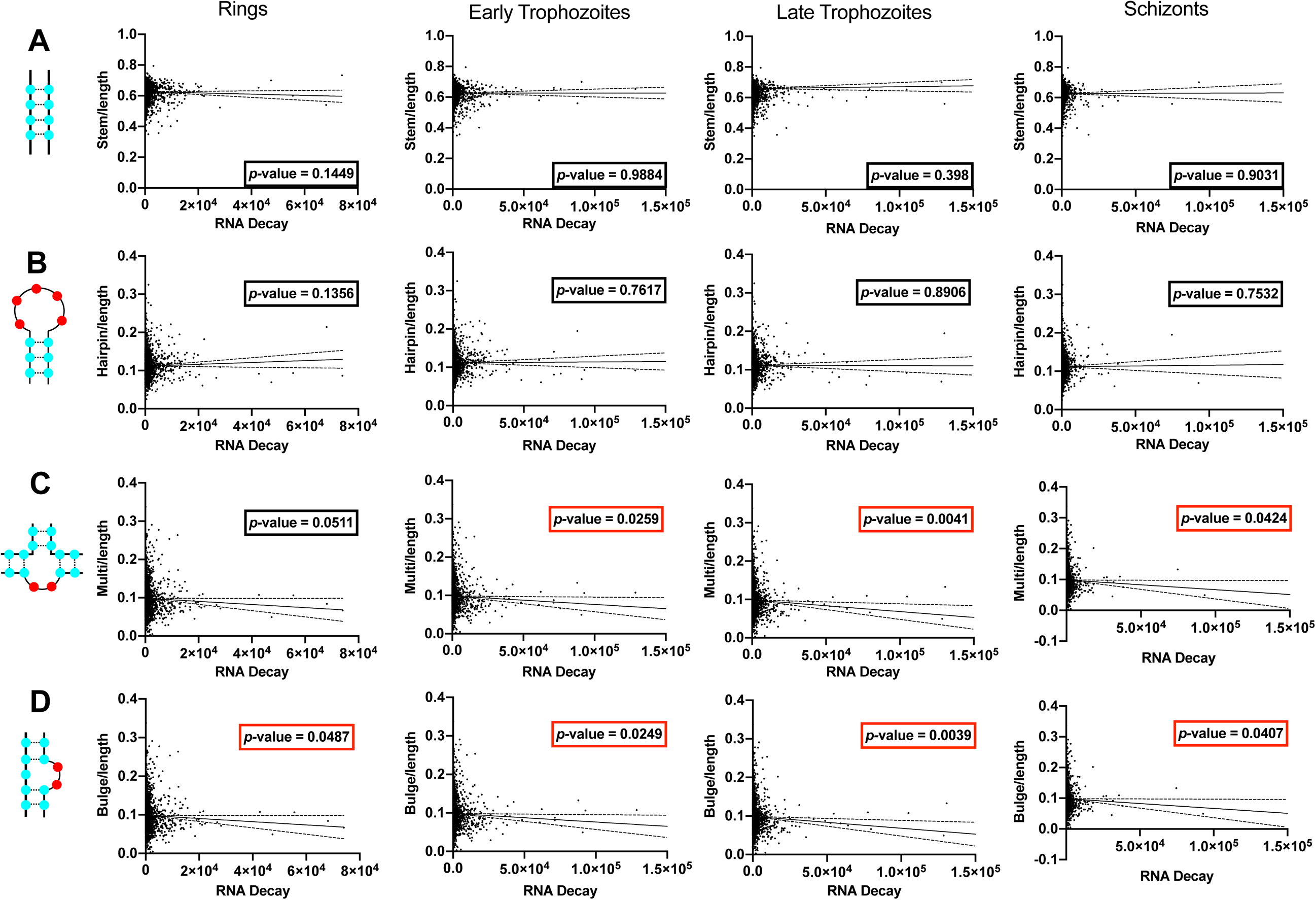
Correlation between RNA secondary structures and RNA decay across the life cycle of *P. falciparum*. The columns represent the different erythrocytic life stages of *P. falciparum* and the rows represent the correlation between RNA decay and **(A)** the number of stems, **(B)** the number of hairpins, **(C)** the number of multi-stem loops and **(D)** the number of bulges. Statistically significant negative correlations have a *p*-value framed in red.

## DISCUSSION

In this study, we produced the SHAPE-seq-determined structurome of a parasitic protozoan, *P. falciparum.* This had never previously been performed for any protozoan parasite, although a second study, appearing after the submission of this manuscript, has also now described an *in vivo P. falciparum* structurome (Qi et al. 2021). This study covered only 23-35% of the transcriptome (in trophozoites and rings respectively), compared to >51% here, and had approximately 35% fewer reads per sample with markedly lower mappability of reads. Our study therefore remains the most comprehensive structurome in *P. falciparum* to date. A third study, equally recently published, mapped duplex RNA regions by RNA-seq after degradation of ssRNA (Alvarez et al. 2021); however this does not produce data directly comparable to an NAI-probed *in vivo* structurome.

Firstly, we demonstrated that it is possible to use probing chemicals on the parasite within the host erythrocyte, meaning that the chemicals can get through the host cell membrane, the parasitophorous vacuole and the parasite membrane. The fact that we could also detect mitochondrial and apicoplast transcripts shows that the chemicals also passed through the inner organellar membranes. Thus, we obtained *in vivo* structures for about half the transcripts in the transcriptome, including structural and noncoding RNAs as well as mRNAs. Most of the structural RNAs had stereotypical structures, although a subset of tRNAs did not model stereotypically – an interesting observation that merits further investigation. Amongst mRNAs, the *in vivo* structures tended to have lower free energy, making them more likely to form, than the equivalent structures predicted *in silico*, thus demonstrating the value of the experimental approach. Highly conserved RNAs such as ribosomal RNAs tended to be predicted well, whereas classes of genes whose structures were poorly predicted *in silico* included genes involved in metabolism, biosynthesis of complex molecules, and transport. These genes may have become particularly divergent in *P. falciparum* due to its parasitic intracellular lifestyle.

We also took advantage of the striking disparities in genome composition in the *Plasmodium* genus to see if the extreme base composition of the *P. falciparum* transcriptome might have an impact on the different structures formed, by comparing the predicted structurome of *P. knowlesi* with that of *P. falciparum*. We observed that in *P. falciparum* transcripts were more prone to form structures like hairpins and multi-stem loops, whereas a more balanced genome like *P. knowlesi* gave rise to orthologous transcripts with more stems. Overall, *P. falciparum* transcripts were predicted to fold with higher free energy than their orthologs in *P. knowlesi*, suggesting that an A/U-biased transcriptome is less stable. However, this might be mitigated by *in vivo* factors, since we also found that overall the *P. falciparum in vivo* structurome had lower free energy than was predicted *in silico*. It would be interesting in future to perform an *in vivo* structurome of *P. knowlesi* and see whether *in vivo* transcripts tend towards greater stability in this species as well.

To examine how the RNA structurome might influence parasite biology, we looked at the impacts of the different structures on translation efficiency and RNA decay. We showed that RNA structures had no impact on translation efficiency in rings and early trophozoites, two stages of the erythrocytic cycle that are translationally less active (Caro et al. 2014; Bunnik et al. 2016). On the contrary, late trophozoites and schizonts are more translationally active and negative correlations between translation efficiency and complex structures like hairpins, multi-loop stem and bulges were observed. Recently, we studied a particularly stable RNA structure in *P. falciparum*, the RNA G-quadruplex, and demonstrated that it likewise had a negative impact on translation (Dumetz et al. 2021). We also observed a positive correlation between translation efficiency and a simpler RNA structure, stems. This is consistent with the idea that stem structures can facilitate ribosome attachment, whereas other complex structures can inhibit it (Kramer and Gregory 2018).

The correlations described above were not detected in either of the parallel *P. falciparum* ‘structurome’ studies. A study comparing duplex RNA regions with translation efficiency failed to observe any clear correlations (Alvarez et al. 2021), probably because it did not distinguish the subtleties of different types of RNA secondary structure: this highlights the power of the structurome approach taken here. Meanwhile, the comparable study by Wang and co-workers (Qi et al. 2021) reported that the least-structured transcripts had the highest abundance, but since abundance was derived only from a snapshot RNA-seq measurement, it was not possible to attribute this specifically to transcript half-life, decay rate, etc. as individual factors that might be affected by RNA structure. Our study examined two such factors individually – translation efficiency and also RNA decay rate – and therefore generated different insights.

There was a negative correlation between structures and RNA decay, so we can hypothesise that RNA structures influence protein expression on at least two levels: the rate of translation and the degree of transcript stability. Similar effects were previously demonstrated in human cell lines, and furthermore RNA base modifications were also found to be important, impacting protein expression by changing the functional half-life of transcripts (Mauger et al. 2019). This latter area has been little explored in *P. falciparum* as yet, but a recent study of base modification in *P. falciparum* likewise implicated RNA base modification in protein expression (Baumgarten et al. 2019).

This study focuses on RNA structure in the erythrocytic life cycle of *Plasmodium* parasites. Two more parts of the full life cycle remain to be studied in future: the liver stage and the sexual cycle which happens in the host vector, a mosquito. The liver stages are particularly interesting because two human-infective *Plasmodium* species, *P. vivax* and *P. ovale*, can make hypnozoites – a dormant form that can stay inside the liver of infected people for months and then reactivate to cause disease (White and Imwong 2012). Currently the biology of hypnozoites remains complicated to investigate. However, translational repression may be key to long-term dormancy, so the RNA structurome of hypnozoites could be intriguing. A more accessible stage, the gametocyte, also shows dormancy (albeit a dormancy that lasts days rather than months), and female gametocytes are known to translationally repress hundreds of transcripts (Guerreiro et al. 2014). Once in the mosquito midgut, the translational repression is lifted, and we can hypothesise that the change of temperature between a warm-blooded host and a non-thermoregulatory insect would impact RNA structures, playing a role in lifting the repression.

Within the mosquito, the sexual life cycle encompasses four different life stages for which no RNA structures are known. This step of the life cycle is crucially important in completion of the transmission chain and also in generating genetic diversity (Arez et al. 2003). Recent progress in single cell RNA sequencing give us the opportunity to look at the transcriptome of single parasites at all life cycle stages, including those in the host vector (Bogale et al. 2021). In parallel, novel RNA structure determination techniques, like SHAPE-MaP-seq associated with the latest algorithms like DRACO, would make the determination of single cell RNA structures possible (Morandi et al. 2021). Other technological breakthroughs like SMRT-seq (Single Molecule, Real Time) associated with SHAPE-MaP would allow access to transcriptional variant structures (Mays et al. 2019), which were not possible to analyse in this study, and this could shed new light on post-transcriptional regulation in *Plasmodium*. Finally, the key roles of RNA-binding proteins in determining RNA structures *in vivo* have yet to be examined. Broadly, Wang and co-workers found that de-proteinised RNAs that had been refolded *in vitro* were less structured, and/or protein-bound, than the same RNAs *in vivo*, which is not unexpected (Qi et al. 2021). However, more specific future experiments could potentially knock out single RNA binding proteins and then compare the resultant RNA structures *in vivo*. This would shed more detailed light on the determinants of the *P. falciparum* RNA structurome.

Overall, this work paves the way to decipher another level of complexity in the molecular biology of *Plasmodium* parasites.

## MATERIAL AND METHODS

### Parasite culture

*Plasmodium falciparum* 3D7, obtained from the Malaria Research and Reference Reagent Resource Center (MR4), was cultured in O+ human erythrocytes at 4% haematocrit in RPMI 1640 supplemented with 0.25 % albumax (Invitrogen), 5% heat-inactivated human serum and 0.25% sodium bicarbonate in gassed chambers at 1% O_2_, 3% CO_2_, and 96% N_2_ (Trager and Jensen 1976). Parasite count was assessed on thin blood smears stained with Hemacolor (Merck).

### RNA structure probing and poly(A) enrichment

An asynchronous culture was grown up to 1.2×10^10^ parasites as previously described (Trager and Jensen 1976). Erythrocytes were pelleted and then incubated for 30 minutes either with 10 mM DMS (dimethyl sulfate, Sigma) in ethanol or 20 mM NAI (2-methylnicotinic acid imidazolide) in anhydrous DMSO, at 37°C, agitated at 200 rpm, in the same gas mixture described above. The NAI was prepared as previously described (Kwok et al. 2013). The reaction was stopped with 40 mM dithiothreitol (DTT). Parasites were extracted by erythrocyte lysis using 1v of 0.2% saponin in PBS and then washed 3 times in ice-cold PBS. RNA was extracted using Qiagen RNeasy plus mini kit (Qiagen) following manufacturer’s instructions and then submitted to DNase treatment for 15 minutes (Qiagen). The total RNA recovered was split into samples of 5 μg each and enriched in poly(A) transcripts using NEBNext Poly(A) mRNA magnetic isolation module (NEB) according to manufacturer’s instructions.

### Visualisation of 5.8S rRNA structure using RT-stalling

RT stalling was performed using a primer targeting the 5.8S rRNA gene (PF3D7_0531800). Briefly, 1.5 μg of total RNA from DMS, NAI and DMSO-treated cultures in 5.5 µl nuclease-free water was mixed with 1 µl of 5 µM Pfa_5.8S primer labelled at the 5’ end with Cy5 [Cy5]ATTTTCTGTAGGAGTACCACT (Eurofins Genomics) in 3 µl of reverse transcription buffer (20 mM Tris, pH 7.5, 4 mM MgCl_2_, 1 mM DTT, 0.5 mM dNTPs, 150 mM LiCl in final). For the sequencing reactions (A/G/C/T), an extra 1 µl of 10 mM dideoxynucleoside triphosphate (Roche) was added to replace 1 µl of nuclease-free water. All samples were heated at 75°C for 3 min, 35°C for 5 min and then held at 50°C. Next, 0.5 µl of Superscript III (200U/µl) (Thermo Scientific) was added and reverse transcription was performed at 50°C for 15 min, followed by addition of 0.5 µl of 2M NaOH at 95°C for 10 min to degrade the RNA template. After reverse transcription, 10 µl of 2x formamide orange G dye (94% deionized formamide, 20 mM Tris pH 7.5, 20 mM EDTA, orange G dye) was added and heated at 95°C for 3 min before loading into a pre-heated (at 90W for 45 min) 8% denaturing polyacrylamide gel. The gel was run at constant power of 90 W for 90 min after loading 3.2 µl of each sample. Fujifilm FLA 9000 was used to scan the gel.

### Library preparation and sequencing details

Random fragmentation of polyA-enriched RNA was performed with 150 ng polyA-enriched RNA in fragmentation buffer (40 mM Tris-HCl pH 8.2, 100 mM LiCl, 30 mM MgCl_2_) at 95°C for 60s to generate average fragment size of ~250 nt, and then purified with RNA Clean & Concentrator (Zymo Research). 3’ dephosphorylation reactions included 7 µl fragmented sample, 1 µl of 10x PNK buffer (NEB), 1 µl of rSAP enzyme (NEB) and 1 µl of PNK enzyme (NEB), carried out at 37°C for 30 min. 3’ adapter ligation was performed by adding 3’ adapter (5’-/5rApp/NNNNNNAGATCGGAAGAGCACACGTCTG/3SpC3/-3’ with 1:5 molar ratio of RNA to 3’ adapter), PEG 8000 (17.5% final) and T4 RNA ligase 2 (NEB) in 1x T4 RNA ligase buffer at 25°C for 1 h. Excess adapter was digested by adding 1 µl RecJf (NEB) and 1 µl 5’deadenylase (NEB) at 30°C for 30 min and removed by RNA Clean & Concentrator (Zymo Research). Reverse transcription reactions including the ligated RNA above, reverse primer (5’-CAGACGTGTGCTCTTCCGATCT-3’ with 1:2.5 molar ratio of RNA to reverse primer), reverse transcription buffer (see RTS method above) and Superscript III (1U/µl final) (Thermo Scientific) were performed at 75°C for 3 min, 35°C for 5 min, then 50°C for 50 min. In this step, Superscript III should be added before the 50°C and after reverse transcription, NaOH (0.1 M final) was added at 95 °C for 10 min for RNA degradation, followed by mixing with 5 µl 1M Tris-HCl (pH 7.5) before RNA Clean & Concentrator (Zymo Research). The purified cDNA was then ligated with 5’ adapter (5’/5Phos/AGATCGGAAGAGCGTCGTGTAGCTCTTCCGATCTN10/3SpC3/-3’ with 1:20 molar ratio of RNA to 5’adapter) using Quick Ligation Kit (NEB) at 37°C overnight. Ligated reaction mixture was heated to 95°C for 10 min for inactivation and mixed with 1 volume of 2x formamide orange G dye before purifying with 10% denaturing urea-TBE acrylamide gel (Thermo Scientific) at 300V for 20 min. The size of 100-400 nt was sliced, crushed and soaked in 1x TEN 250 buffer (1x TE pH 7.4, 0.25 M NaCl) with incubation at 80°C for 30 min with 1300 rpm shaking, and then purified with RNA Clean & Concentrator (Zymo Research). The purified ssDNA was mixed with forward primer (5’-AATGATACGGCGACCACCGAGATCTACACTCTTTCCCTACACGACGCTCTTCCG ATCT-3’, 0.5 µM final) and reverse primer with different indexes (5’-CAAGCAGAAGACGGCATACGAGAT-(6 nt index seq)-GTGACTGGAGTTCAGACGTGTGCTCTTCCGATCT-3’, 0.5 µM final) for PCR amplification by using KAPA HiFi HotStart ReadyMix (KAPA Biosystems). The PCR program included 95°C for 3 min, 16 cycles of 3 steps (98°C: 20s, 68°C: 15s, 72°C: 40s), 72°C for 90s, then cooled to 4°C for size selection. PCR samples were resolved on a 1.8% TAE agarose gel at 120 V for 55 min. The size of 150-400 nt was cut and recovered by Zymo DNA agarose gel extraction kit (Zymo Research). DNA libraries were quantified, pooled and sequenced on the Illumina Hiseq System in 150 bp paired-end (PE) configuration. A more detail protocol was described in reference (Yeung et al. 2019).

### Structurome analysis

Forward and reverse reads from each sample were cleaned of adapter sequence contamination using *reaper* (v15-065) from the *kraken* package (Davis et al. 2013). Paired reads were then aligned to an indexed reference transcriptome (PlasmoDB-45_Pfalciparum3D7) using *Bowtie2* (v2.4.2) in paired-end mode and converted to SAM and sorted BAM files (Langmead et al. 2019). Reverse transcription stops were assessed using *StructureFold2* for each individual sample (Tack et al. 2018).

Coverage and overlap of RT stop data were then computed along with cross-replicate coverage and overlap and stop correlation was assessed among all samples via *StructureFold2*. These data were normalised and structural reactivity data generated for each transcript with enough supporting data across samples using *StructureFold2*. These reactivities were supplied to *RNAStructure* for experimental reactivity constrained folding (Reuter and Mathews 2010). Resulting structures were then assessed for structural features using *Forgi* from the *ViennaRNA* package (Lorenz et al. 2011). Unconstrained folding of every transcript was also performed using RNAStructure with the same parameters and assessed similarly to generate a cohort of unconstrained purely *in silico* folded transcripts.

Parameters for the number of reads and mapping efficiency for each library are shown (Table 1).

**Table 1:** Sequencing parameters for each NGS library generated in this study.

| Sample | Number of<br>Read Pairs | Yield<br>(Gigabases) | Mapped<br>Reads | Mapped<br>Reads (%) |
| --- | --- | --- | --- | --- |
| Total-RNA-DMS-3 | 148,865,508 | 44.6597 | 108,912,838 | 73.1% |
| Total-RNA-DMSO-3 | 163,681,632 | 49.1045 | 115,215,035 | 70.3% |
| Total-RNA-NAI-3 | 192,424,134 | 57.7272 | 144,128,728 | 74.9% |
| Total-RNA-DMS-2 | 166,742,223 | 50.0227 | 117,106,285 | 70.2% |
| Total-RNA-DMSO-2 | 186,503,272 | 55.951 | 128,887,798 | 69.1% |
| Total-RNA-NAI-2 | 145,601,085 | 43.6803 | 108,451,225 | 74.5% |

### Functional dataset analysis

Microarray datasets from parasites exposed to hyperoxia were extracted from (Torrentino-Madamet et al. 2011), and the chloroquine-exposed dataset was extracted from (Untaroiu et al. 2019). Lists of genes were updated to the current genome annotation. Ribosome profiling data were obtained from (Caro et al. 2014) and translational efficiency was calculated as ribosome density per messenger RNA. The RNA decay dataset was extracted from (Painter et al. 2018) at 10hpi for rings, 24hpi for early trophozoites, 30hpi for late trophozoites and 46hpi for schizonts. Structural parameters per transcript (numbers of nucleotides involved in stems, hairpins, multi-stem loop and bulges) were normalized to transcript length before comparing these structural parameters with the translational efficiency of each gene.

## Acknowledgements

We would like to thank Dr. Betty Chung from the Cambridge Pathology Department for her help with the transcription efficiency data processing.

## Authors’ contributions

FD performed experiments, data analysis, made the figures and wrote the manuscript, AE performed the bioinformatic Structure-seq analysis, JZ performed the structure probing gel assay and Structure-seq library preparation and sequencing, and participated in the writing, CKK and CJM conceptualised the paper and wrote the manuscript.

## REFERENCES

Alvarez DR, Ospina A, Barwell T, Zheng B, Dey A, Li C, Basu S, Shi X, Kadri S, Chakrabarti K. 2021. The RNA structurome in the asexual blood stages of malaria pathogen plasmodium falciparum. RNA Biol 00: 1–18. https://doi.org/10.1080/15476286.2021.1926747.

Arez AP, Pinto J, Pålsson K, Snounou G, Jaenson TGT, do Rosário VE. 2003. Transmission of mixed Plasmodium species and Plasmodium falciparum genotypes. Am J Trop Med Hyg 68: 161–8. http://www.ncbi.nlm.nih.gov/pubmed/12641406.

Aurrecoechea C, Brestelli J, Brunk BP, Dommer J, Fischer S, Gajria B, Gao X, Gingle A, Grant G, Harb OS, et al. 2009. PlasmoDB: a functional genomic database for malaria parasites. Nucleic Acids Res 37: D539–D543. http://www.ncbi.nlm.nih.gov/pubmed/18957442.

Baumgarten S, Bryant JM, Sinha A, Reyser T, Preiser PR, Dedon PC, Scherf A. 2019. Transcriptome-wide dynamics of extensive m6A mRNA methylation during Plasmodium falciparum blood-stage development. Nat Microbiol 4: 2246–2259. http://dx.doi.org/10.1038/s41564-019-0521-7.

Bogale HN, Pascini T V., Kanatani S, Sá JM, Wellems TE, Sinnis P, Vega-Rodríguez J, Serre D. 2021. Transcriptional heterogeneity and tightly regulated changes in gene expression during Plasmodium berghei sporozoite development. Proc Natl Acad Sci 118: e2023438118. http://www.pnas.org/lookup/doi/10.1073/pnas.2023438118.

Bunnik EM, Batugedara G, Saraf A, Prudhomme J, Florens L, Le Roch KG. 2016. The mRNA-bound proteome of the human malaria parasite Plasmodium falciparum. Genome Biol 17: 1–18. http://dx.doi.org/10.1186/s13059-016-1014-0.

Caro F, Ahyong V, Betegon M, DeRisi JL. 2014. Genome-wide regulatory dynamics of translation in the Plasmodium falciparum asexual blood stages. Elife 3: 1–24. https://elifesciences.org/articles/04106.

Casas-Vila N, Sayols S, Pérez-Martínez L, Scheibe M, Butter F. 2020. The RNA fold interactome of evolutionary conserved RNA structures in S. cerevisiae. Nat Commun 11: 1–12. http://dx.doi.org/10.1038/s41467-020-16555-4.

Cech TR, Steitz JA. 2014. The noncoding RNA revolution-trashing old rules to forge new ones. Cell 157: 77–94. http://dx.doi.org/10.1016/j.cell.2014.03.008.

Davis MPA, van Dongen S, Abreu-Goodger C, Bartonicek N, Enright AJ. 2013. Kraken: A set of tools for quality control and analysis of high-throughput sequence data. Methods 63: 41–49. http://dx.doi.org/10.1016/j.ymeth.2013.06.027.

Deng H, Cheema J, Zhang H, Woolfenden H, Norris M, Liu Z, Liu Q, Yang X, Yang M, Deng X, et al. 2018. Rice In Vivo RNA Structurome Reveals RNA Secondary Structure Conservation and Divergence in Plants. Mol Plant 11: 607–622. https://doi.org/10.1016/j.molp.2018.01.008.

Ding Y, Tang Y, Kwok CK, Zhang Y, Bevilacqua PC, Assmann SM. 2014. In vivo genome-wide profiling of RNA secondary structure reveals novel regulatory features. Nature 505: 696–700. http://dx.doi.org/10.1038/nature12756.

Dumetz F, Chow EY, Harris LM, Umar MI, Jensen A, Chung B, Chan TF, Merrick CJ, Kwok CK. 2021. G-quadruplex RNA motifs influence gene expression in the malaria parasite Plasmodium falciparum. Nucleic Acids Research 1 doi.org/10.1093/nar/gkab1095.

Fang J, McCutchan TF. 2002. Thermoregulation in a parasite’s life cycle. Nature 418: 742.

Fang J, Zhou H, Rathore D, Sullivan M, Su XZ, McCutchan TF. 2004. Ambient glucose concentration and gene expression in Plasmodium falciparum. Mol Biochem Parasitol 133: 125–129.

Gardner MJ, Hall N, Fung E, White O, Berriman M, Hyman RW, Carlton JM, Pain A, Nelson KE, Bowman S, et al. 2002. Genome sequence of the human malaria parasite Plasmodium falciparum. Nature 419: 498–511.

Guerreiro A, Deligianni E, Santos JM, Silva PAGC, Louis C, Pain A, Janse CJ, Franke-Fayard B, Carret CK, Siden-Kiamos I, et al. 2014. Genome-wide RIP-Chip analysis of translational repressor-bound mRNAs in the Plasmodium gametocyte. Genome Biol 15: 1–16.

Gunderson JH, Sogin ML, Wollett G, Hollingdale M, de la Cruz VF, Waters AP, McCutchan TF. 1987. Structurally distinct, stage-specific ribosomes occur in Plasmodium. Science 238: 933–7. http://www.ncbi.nlm.nih.gov/pubmed/3672135.

Guo JU, Bartel DP. 2016. RNA G-quadruplexes are globally unfolded in eukaryotic cells and depleted in bacteria. Science (80-) 353.

Hasler D, Meister G. 2016. From tRNA to miRNA: RNA-folding contributes to correct entry into noncoding RNA pathways. FEBS Lett 590: 2354–63. http://www.ncbi.nlm.nih.gov/pubmed/27397696.

Huber RG, Lim XN, Ng WC, Sim AYL, Poh HX, Shen Y, Lim SY, Sundstrom KB, Sun X, Aw JG, et al. 2019. Structure mapping of dengue and Zika viruses reveals functional long-range interactions. Nat Commun 10. http://dx.doi.org/10.1038/s41467-019-09391-8.

Huston NC, Wan H, Strine MS, de Cesaris Araujo Tavares R, Wilen CB, Pyle AM. 2021. Comprehensive in vivo secondary structure of the SARS-CoV-2 genome reveals novel regulatory motifs and mechanisms. Mol Cell 81: 584–598.e5. https://linkinghub.elsevier.com/retrieve/pii/S109727652030962X.

Incarnato D, Neri F, Anselmi F, Oliviero S. 2014. Genome-wide profiling of mouse RNA secondary structures reveals key features of the mammalian transcriptome. Genome Biol 15: 491. http://www.ncbi.nlm.nih.gov/pubmed/25323333.

John Rogers M, Gutell RR, Damberger SH, Li J, Mcconkey GA, Waters AP, McCutchan TF. 1996. Structural features of the large subunit rRNA expressed in Plasmodium falciparum sporozoites that distinguish it from the asexually expressed large subunit rRNA. RNA 2: 134–145.

Kramer MC, Gregory BD. 2018. Does RNA secondary structure drive translation or vice versa? Nat Struct Mol Biol 25: 641–643.

Kwok CK, Ding Y, Tang Y, Assmann SM, Bevilacqua PC. 2013. Determination of in vivo RNA structure in low-abundance transcripts. Nat Commun 4: 2971. http://www.nature.com/articles/ncomms3971.

Kwok CK, Tang Y, Assmann SM, Bevilacqua PC. 2015. The RNA structurome: transcriptome-wide structure probing with next-generation sequencing. Trends Biochem Sci 40: 221–32. http://dx.doi.org/10.1016/j.tibs.2015.02.005.

Langmead B, Wilks C, Antonescu V, Charles R. 2019. Scaling read aligners to hundreds of threads on general-purpose processors ed. J. Hancock. Bioinformatics 35: 421–432. http://www.ncbi.nlm.nih.gov/pubmed/30020410.

Li P, Wei Y, Mei M, Tang L, Sun L, Huang W, Zhou J, Zou C, Zhang S, Qin CF, et al. 2018. Integrative Analysis of Zika Virus Genome RNA Structure Reveals Critical Determinants of Viral Infectivity. Cell Host Microbe 24: 875–886.e5. https://doi.org/10.1016/j.chom.2018.10.011.

Ling Yang S, DeFalco L, Anderson DE, Zhang Y, Aw AJ, Lim Y, Xin Ni L, Yee Tan K, Zhang T, Chawla T, et al. 2021. Comprehensive mapping of SARS-CoV-2 interactions in vivo reveals 1 functional virus-host interactions 2 3. bioRxiv 2021.01.17.427000. https://doi.org/10.1101/2021.01.17.427000.

Lorenz R, Bernhart SH, Höner Zu Siederdissen C, Tafer H, Flamm C, Stadler PF, Hofacker IL. 2011. ViennaRNA Package 2.0. Algorithms Mol Biol 6: 26. http://www.ncbi.nlm.nih.gov/pubmed/22115189.

Manfredonia I, Nithin C, Ponce-Salvatierra A, Ghosh P, Wirecki TK, Marinus T, Ogando NS, Snijder EJ, van Hemert MJ, Bujnicki JM, et al. 2020. Genome-wide mapping of SARS-CoV-2 RNA structures identifies therapeutically-relevant elements. Nucleic Acids Res 1–17. https://academic.oup.com/nar/advance-article/doi/10.1093/nar/gkaa1053/5961787.

Mathews DH, Sabina J, Zuker M, Turner DH. 1999. Expanded sequence dependence of thermodynamic parameters improves prediction of RNA secondary structure. J Mol Biol 288: 911–40. http://www.ncbi.nlm.nih.gov/pubmed/10329189.

Mauger DM, Cabral BJ, Presnyak V, Su S V., Reid DW, Goodman B, Link K, Khatwani N, Reynders J, Moore MJ, et al. 2019. mRNA structure regulates protein expression through changes in functional half-life. Proc Natl Acad Sci U S A 116: 24075–24083. http://www.pnas.org/lookup/doi/10.1073/pnas.1908052116.

Mays AD, Schmidt M, Graham G, Tseng E, Baybayan P, Sebra R, Sanda M, Mazarati JB, Riegel A, Wellstein A. 2019. Single-molecule real-time (SMRT) full-length RNA-sequencing reveals novel and distinct mRNA isoforms in human bone marrow cell subpopulations. Genes (Basel) 10: 1–17.

Morandi E, Manfredonia I, Simon LM, Anselmi F, van Hemert MJ, Oliviero S, Incarnato D. 2021. Genome-scale deconvolution of RNA structure ensembles. Nat Methods 18: 249–252. http://www.ncbi.nlm.nih.gov/pubmed/33619392.

Pain A, Böhme U, Berry AE, Mungall K, Finn RD, Jackson AP, Mourier T, Mistry J, Pasini EM, Aslett MA, et al. 2008. The genome of the simian and human malaria parasite Plasmodium knowlesi. Nature 455: 799–803.

Painter HJ, Chung NC, Sebastian A, Albert I, Storey JD, Llinás M. 2018. Genome-wide real-time in vivo transcriptional dynamics during Plasmodium falciparum blood-stage development. Nat Commun 9: 2656. http://dx.doi.org/10.1038/s41467-018-04966-3.

Qi Y, Zhang Y, Zheng G, Chen B, Zhang M, Li J, Peng T, Huang J, Wang X. 2021. In Vivo and In Vitro Genome-Wide Profiling of RNA Secondary Structures Reveals Key Regulatory Features in Plasmodium falciparum. Front Cell Infect Microbiol 11: 1–13. https://www.frontiersin.org/articles/10.3389/fcimb.2021.673966/full.

Reuter JS, Mathews DH. 2010. RNAstructure: software for RNA secondary structure prediction and analysis. BMC Bioinformatics 11: 129. https://bmcbioinformatics.biomedcentral.com/articles/10.1186/1471-2105-11-129.

Righetti F, Nuss AM, Twittenhoff C, Beele S, Urban K, Will S, Bernhart SH, Stadler PF, Dersch P, Narberhaus F. 2016. Temperature-responsive in vitro RNA structurome of Yersinia pseudotuberculosis. Proc Natl Acad Sci U S A 113: 7237–7242.

Ritchey LE, Su Z, Tang Y, Tack DC, Assmann SM, Bevilacqua PC. 2017. Structure-seq2: sensitive and accurate genome-wide profiling of RNA structure in vivo. Nucleic Acids Res 45: e135.

Rouskin S, Zubradt M, Washietl S, Kellis M, Weissman JS. 2014. Genome-wide probing of RNA structure reveals active unfolding of mRNA structures in vivo. Nature 505: 701–705. http://dx.doi.org/10.1038/nature12894.

Sharp PA. 2009. The Centrality of RNA. Cell 136: 577–580.

Simon LM, Morandi E, Luganini A, Gribaudo G, Martinez-Sobrido L, Turner DH, Oliviero S, Incarnato D. 2019. In vivo analysis of influenza A mRNA secondary structures identifies critical regulatory motifs. Nucleic Acids Res 47: 7003–7017.

Spitale RC, Crisalli P, Flynn RA, Torre EA, Kool ET, Chang HY. 2013. RNA SHAPE analysis in living cells. Nat Chem Biol 9: 18–20. http://www.ncbi.nlm.nih.gov/pubmed/23178934.

Su Z, Tang Y, Ritchey LE, Tack DC, Zhu M, Bevilacqua PC, Assmann SM. 2018. Genome-wide RNA structurome reprogramming by acute heat shock globally regulates mRNA abundance. Proc Natl Acad Sci U S A 115: 12170–12175.

Sun L, Fazal FM, Li P, Broughton JP, Lee B, Tang L, Huang W, Kool ET, Chang HY, Zhang QC. 2019. RNA structure maps across mammalian cellular compartments. Nat Struct Mol Biol 26: 322–330. http://dx.doi.org/10.1038/s41594-019-0200-7.

Sun L, Li P, Ju X, Rao J, Huang W, Ren L, Zhang S, Xiong T, Xu K, Zhou X, et al. 2021. In vivo structural characterization of the SARS-CoV-2 RNA genome identifies host proteins vulnerable to repurposed drugs. Cell 184: 1865–1883.e20. https://doi.org/10.1016/j.cell.2021.02.008.

Supek F, Bošnjak M, Škunca N, Šmuc T. 2011. REVIGO Summarizes and Visualizes Long Lists of Gene Ontology Terms ed. C. Gibas. PLoS One 6: e21800. http://www.ncbi.nlm.nih.gov/pubmed/21789182.

Tack DC, Tang Y, Ritchey LE, Assmann SM, Bevilacqua PC. 2018. StructureFold2: Bringing chemical probing data into the computational fold of RNA structural analysis. Methods 143: 12–15. https://doi.org/10.1016/j.ymeth.2018.01.018.

Tammy C. T. Lan, Matthew F. Allan, Lauren E. Malsick, Stuti Khandwala, Sherry S. Y. Nyeo, Yu Sun, Junjie U. Guo, Mark Bathe, Anthony Griffiths SR, Lan TCT, Allan MF, Malsick LE, Khandwala S, Nyeo SSY, Sun Y, Guo JU, Bathe M, Griffiths A, et al. 2021. Insights into the secondary structural ensembles of the full SARS-CoV-2 RNA genome in infected cells. bioRxiv 2020.06.29.178343. http://biorxiv.org/content/early/2021/02/19/2020.06.29.178343.abstract.

Tomezsko PJ, Corbin VDA, Gupta P, Swaminathan H, Glasgow M, Persad S, Edwards MD, Mcintosh L, Papenfuss AT, Emery A, et al. 2020. Determination of RNA structural diversity and its role in HIV-1 RNA splicing. Nature 582: 438– 442. http://dx.doi.org/10.1038/s41586-020-2253-5.

Torrentino-Madamet M, Alméras L, Desplans J, Priol Y Le, Belghazi M, Pophillat M, Fourquet P, Jammes Y, Parzy D. 2011. Global response of Plasmodium falciparum to hyperoxia: A combined transcriptomic and proteomic approach. Malar J 10: 1–15.

Trager W, Jensen JB. 1976. Human malaria parasites in continuous culture. Science 193: 673–5. http://www.sciencemag.org/cgi/doi/10.1126/science.781840.

Trotta E. 2014. On the normalization of the minimum free energy of RNAs by sequence length. PLoS One 9.

Twittenhoff C, Brandenburg VB, Righetti F, Nuss AM, Mosig A, Dersch P, Narberhaus F. 2020. Lead-seq: Transcriptome-wide structure probing in vivo using lead(II) ions. Nucleic Acids Res 48: E71–E71.

Untaroiu AM, Carey MA, Guler JL, Papin JA. 2019. Leveraging the effects of chloroquine on resistant malaria parasites for combination therapies. BMC Bioinformatics 20: 1–9.

Vandivier LE, Anderson SJ, Foley SW, Gregory BD. 2016. The Conservation and Function of RNA Secondary Structure in Plants. Annu Rev Plant Biol 67: 463–488.

Velichutina I V., Rogers MJ, McCutchan TF, Liebman SW. 1998. Chimeric rRNAs containing the GTPase centers of the developmentally regulated ribosomal rRNAs of Plasmodium falciparum are functionally distinct. RNA 4: 594–602.

Wachter A. 2010. Riboswitch-mediated control of gene expression in eukaryotes. RNA Biol 7.

Wan Y, Kertesz M, Spitale RC, Segal E, Chang HY. 2011. Understanding the transcriptome through RNA structure. Nat Rev Genet 12: 641–655.

Waters AP, Syin C, McCutchan TF. 1989. Developmental regulation of stage-specific ribosome populations in Plasmodium. Nature 342: 438–40. http://www.ncbi.nlm.nih.gov/pubmed/2586613.

Watts JM, Dang KK, Gorelick RJ, Leonard CW, Bess JW, Swanstrom R, Burch CL, Weeks KM. 2009. Architecture and secondary structure of an entire HIV-1 RNA genome. Nature 460: 711–6. http://www.nature.com/articles/nature08237.

Westermann AJ, Gorski SA, Vogel J. 2012. Dual RNA-seq of pathogen and host. Nat Rev Microbiol 10: 618–630. http://www.ncbi.nlm.nih.gov/pubmed/22890146.

Westhof E, Romby P. 2010. The RNA structurome: high-throughput probing. Nat Methods 7: 965–967. http://dx.doi.org/10.1038/nmeth1210-965.

White NJ, Imwong M. 2012. Relapse. In Advances in Parasitology, pp. 113–150.

WHO. 2018. *World malaria report* 2018. https://www.who.int/malaria/publications/world-malaria-report-2018/en/.

Wilkinson KA, Merino EJ, Weeks KM. 2006. Selective 2’-hydroxyl acylation analyzed by primer extension (SHAPE): quantitative RNA structure analysis at single nucleotide resolution. Nat Protoc 1: 1610–6. http://www.nature.com/articles/nprot.2006.249.

Wu X, Bartel DP. 2017. Widespread Influence of 3′-End Structures on Mammalian mRNA Processing and Stability. Cell 169: 905–917.e11. http://dx.doi.org/10.1016/j.cell.2017.04.036.

Yeung PY, Zhao J, Chow EYC, Mou X, Hong HQ, Chen L, Chan TF, Kwok CK. 2019. Systematic evaluation and optimization of the experimental steps in RNA G-quadruplex structure sequencing. Sci Rep 9: 1–12.

Zhao J, Qian X, Yeung PY, Zhang QC, Kwok CK. 2019. Mapping In Vivo RNA Structures and Interactions. Trends Biochem Sci 44: 555–556. https://doi.org/10.1016/j.tibs.2019.01.012.

Ziv O, Price J, Shalamova L, Kamenova T, Goodfellow I, Weber F, Miska EA. 2020. The Short- and Long-Range RNA-RNA Interactome of SARS-CoV-2. Mol Cell 80: 1067–1077.e5. https://doi.org/10.1016/j.molcel.2020.11.004.

