## Supplementary material for "The *in vivo* RNA structurome of the malaria parasite *Plasmodium falciparum*, a protozoan with an A/T-rich transcriptome": Supp figs and legends

#### Supplementary materials

**Supplemental figure 1:** Polyacrylamide gel showing the reverse-transcribed 5.8S rRNA gene of *P. falciparum* after probing with DMS (2 concentrations), NAI, vehicle control, i.e. DMSO, or no treatment (UT). DMS and NAI modifications are detected as reverse-transcriptase stops, with very similar patterns at both DMS concentrations. The vehicle control confirms that there is negligible 'background', i.e. almost no reverse-transcriptase stops, yielding a near-empty lane similar to the untreated control.

**Supplemental figure 2:** DMS and NAI samples: correlation and pooling data.
(A) For each of the different treatments (both individual replicates and pooled), the distribution of 'RNAdistance' is plotted, i.e. the computed distances between the purely computationally-determined structures and those guided by reactivity data. The y-axis represents the frequency of RNAdistance metrics, which are represented on the x-axis. Replicates of each treatment appear similar, as quantified in (B). (RNAdistance calculated as per: R. Lorenz, S.H. Bernhart, C. Hoener zu Siederdissen, H. Tafer, C. Flamm, P.F. Stadler and I.L. Hofacker (2011), "ViennaRNA Package 2.0", Algorithms for Molecular Biology: 6:26)
B) Pearson correlation heat map showing how the RNAdistance-derived distributions in panel A correlate across samples. Two distinct groupings are clearly formed for NAI and DMS.

**Supplemental figure 3:** Plots showing the coverage of each transcript (averaged per nucleotide) in each replicate of each treatment condition. More than 50% of transcripts in every condition met the coverage threshold of >1.0 per nucleotide.

**Supplemental figure 4:** *RNAStructure* folding of three non-canonical-shaped tRNAs. (A) valine tRNA (PF3D7_0312600), (B) serine tRNA (PF3D7_0410100) and (C) leucine (PF3D7_0620900). The structures in (A) and (C) could also be folded into a less-energetically favourable canonical shape: to exemplify this, NAI-reactive bases in (A) are marked on both structures (red, orange, green from highest to lowest relative reactivity). The structure in B could not be folded into a canonical shape, even at 50% less favourable energy, when using the constraints obtained by NAI probing.

**Supplemental figure 5:** The canonical structure of the *P. falciparum* 18S rRNA (from https://crw-site.chemistry.gatech.edu/) was compared with base reactivity data for the assembled sequence of the PF3D7_0725600 gene (encoding blood-stage-expressed 18S rRNA). Maximum base reactivities for both the NAI and DMS datasets were mapped: NAI reactivities in the blue channel and DMS reactivities in the red channel, hence dually-reactive bases appear pink. Reactivities were scaled to colour intensity.

**Supplemental table 1:** Complete and detailed list of transcripts found during the Structure-seq analysis. The file contains all the genomic information, product description, gene type, transcript length, GC and AT content, reactivity for each replicated of DMS and NAI experiment plus the combined one, and the gene ontology information.

**Supplemental table 2:** Table of transcript variants. Same information available as in Supplementary table 1.

**Supplemental table 3:** REVIGO output for clustering of GO terms. Each tab represents the data per tier of divergence.

**Supplemental table 4:** Table of free energy and transcript structures from Structure-seq and *in silico* analysis.

**Supplemental table 5:** List of the *P. falciparum* transcripts and their *P. knowlesi* orthologs with *in silico* folding information normalised by transcript length.

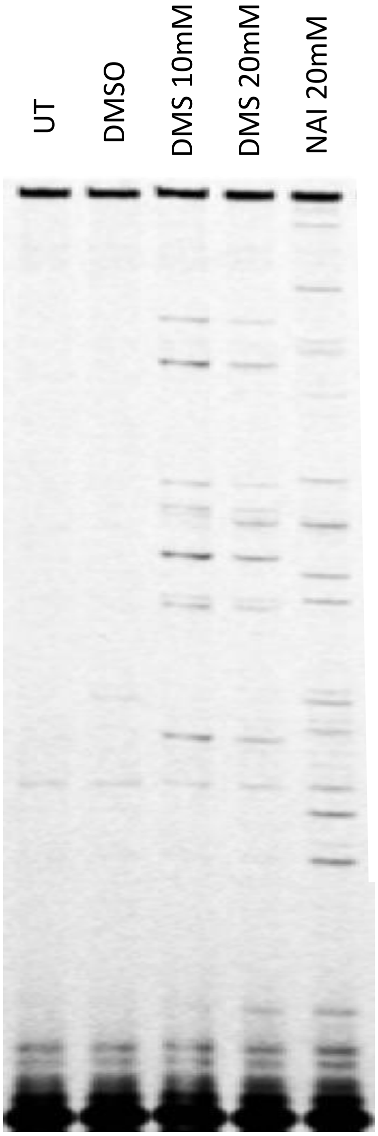


Supplementary figure 1

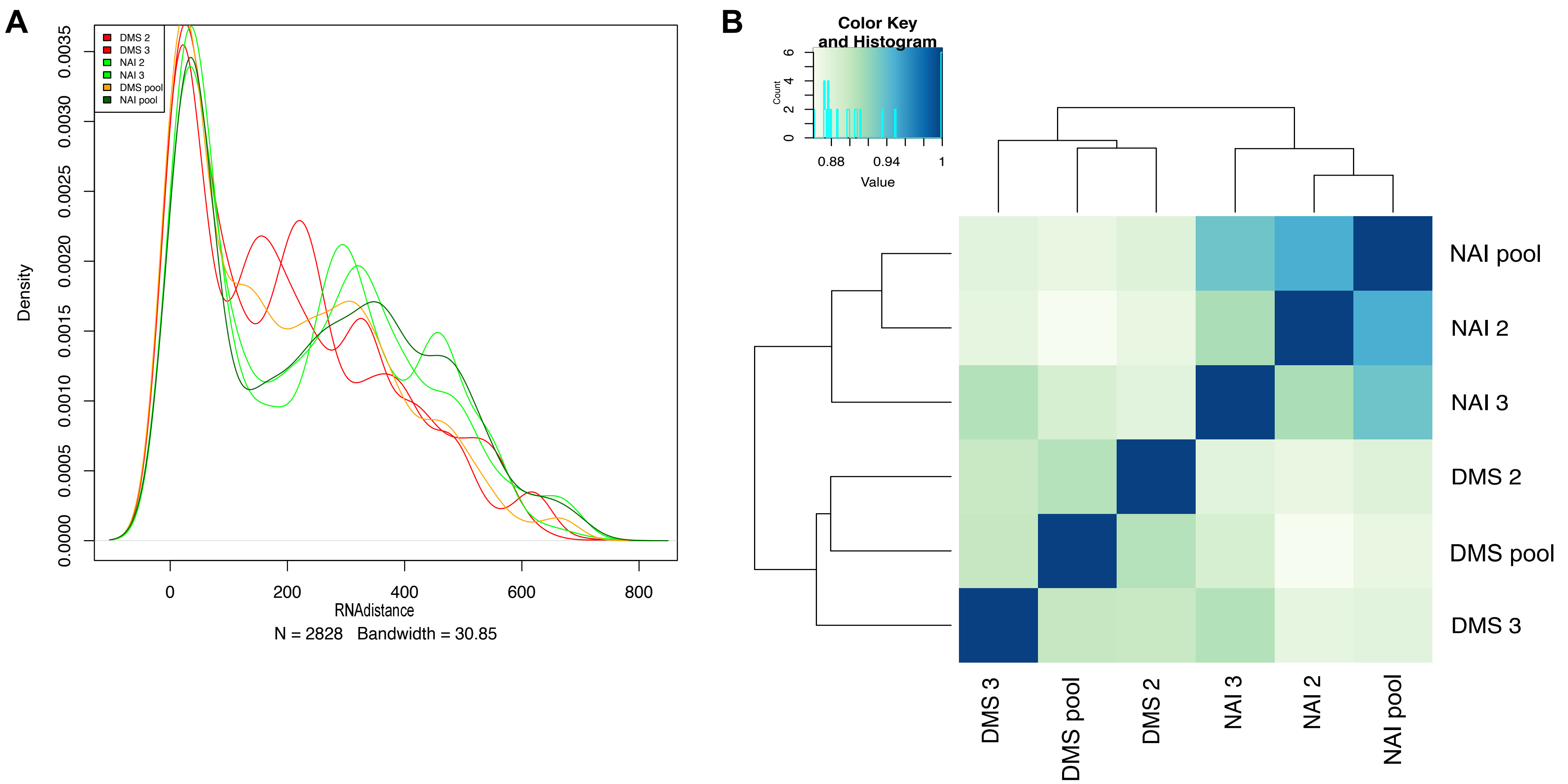


Supplementary figure 2

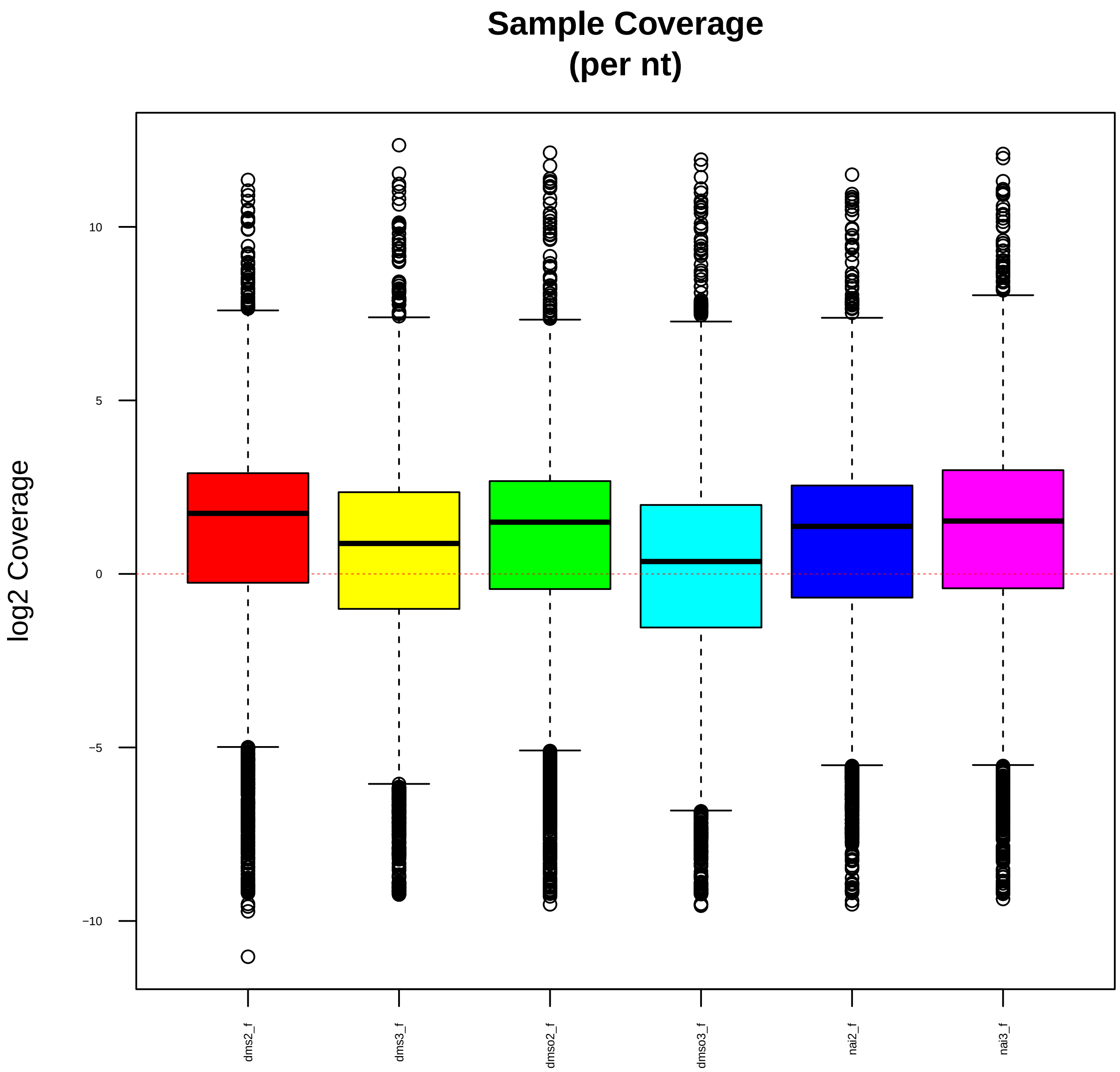


Supplementary figure 3

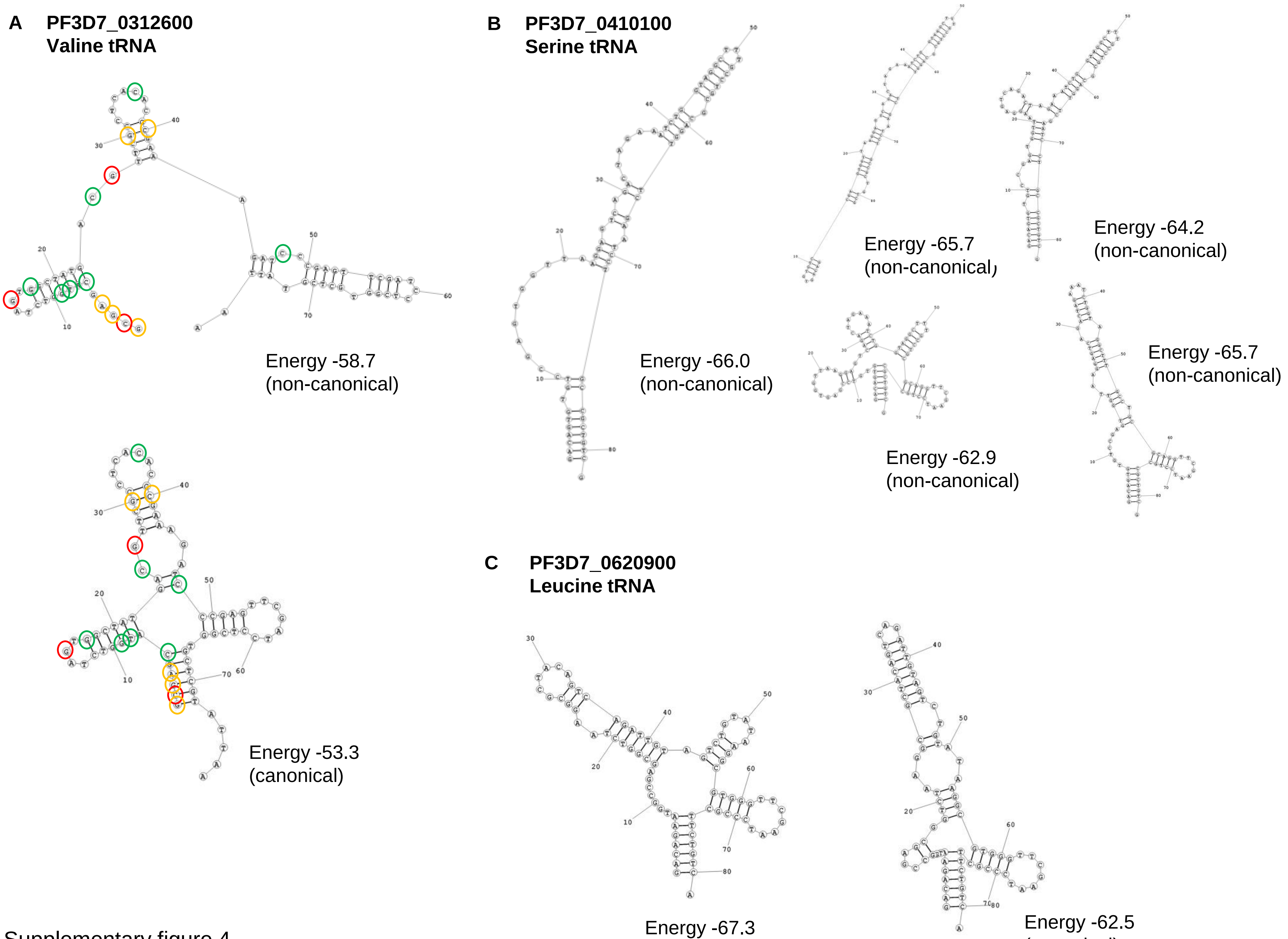


Supplementary figure 4

### Secondary Structure: small subunit ribosomal RNA

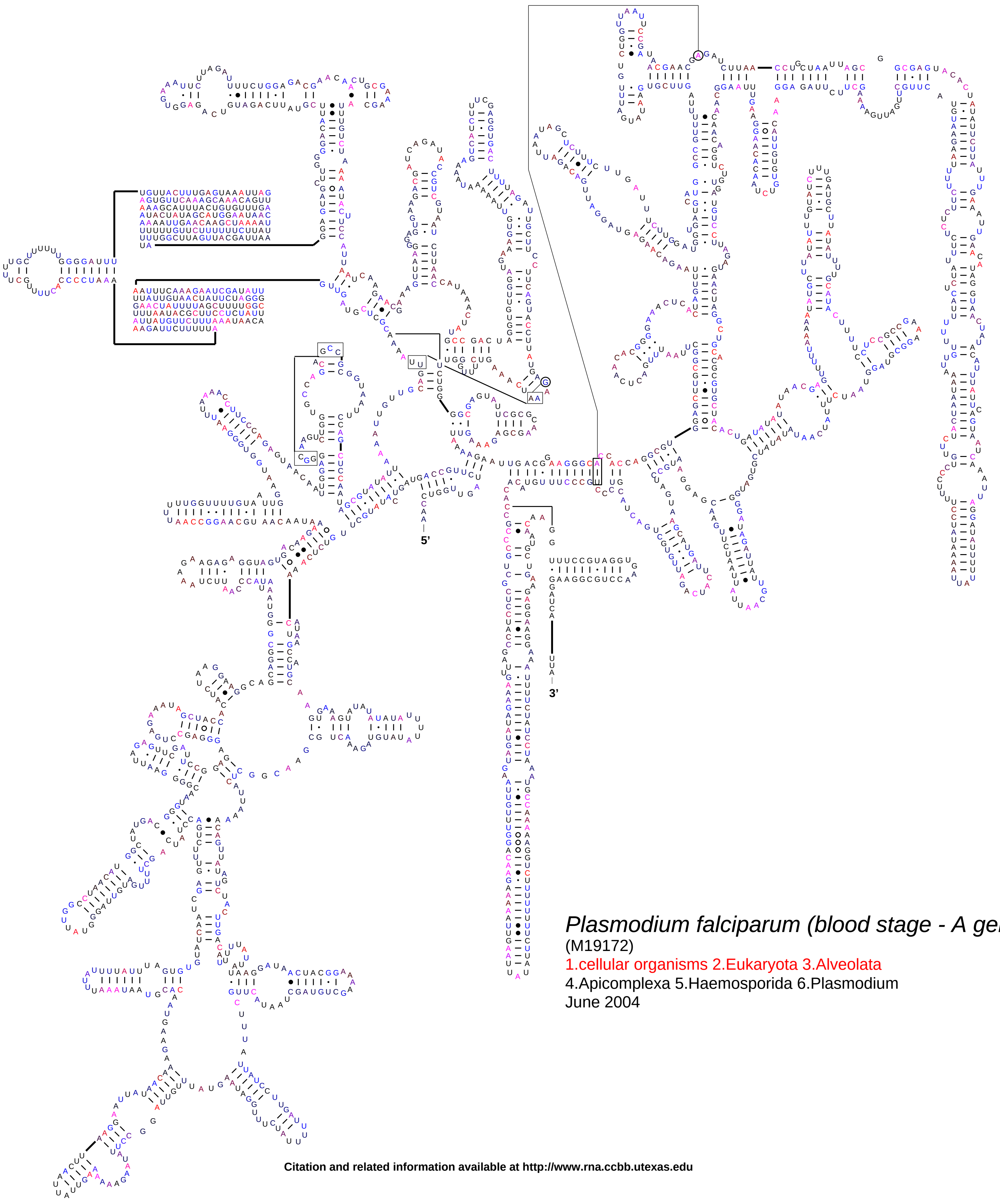


*Plasmodium falciparum (blood stage - A gene)*
(M19172)
1.cellular organisms 2.Eukaryota 3.Alveolata
4.Apicomplexa 5.Haemosporida 6.Plasmodium
June 2004

**Citation and related information available at http://www.rna.ccbb.utexas.edu**
